# A neuromechanical model for *Drosophila* larval crawling based on physical measurements

**DOI:** 10.1101/2020.07.17.208611

**Authors:** Xiyang Sun, Yingtao Liu, Chang Liu, Koichi Mayumi, Kohzo Ito, Akinao Nose, Hiroshi Kohsaka

## Abstract

Animal locomotion requires dynamic interactions between neural circuits, muscles, and surrounding environments. In contrast to intensive studies on neural circuits, the neuromechanical basis for animal behaviour remains unclear due to the lack of information on the physical properties of animals. Here, we proposed an integrated neuromechanical model based on physical measurements by taking *Drosophila* larvae as a model of soft-bodied animals. The biomechanical parameters of fly larvae were measured by the stress-relaxation test. By optimizing parameters in the neural circuit, our neuromechanical model succeeded in quantitatively reproducing the kinematics of larval locomotion that were obtained experimentally. This model could reproduce the observation of optogenetic studies reported previously. The model predicted that peristaltic locomotion could be exhibited in a low friction condition. Analysis of floating larvae provided results consistent with this prediction. Furthermore, the model predicted a significant contribution of intersegmental connections in the central nervous system, which contrasts with a previous study. This hypothesis allowed us to make a testable prediction for the variability in intersegmental connection in sister species of the genus *Drosophila*. Our model based on physical measurement provides a new foundation to study locomotion in soft-bodied animals and soft robot engineering.

## Introduction

Animal behaviour requires both neural circuits and the body. Specific neural circuits referred to as central pattern generators (CPGs) are capable of creating spatiotemporal activity patterns for behaviours (1–5). CPGs regulate motor neuron activity and muscular contraction to generate force for movement. On the other hand, the body’s physical properties and constraints are also significant factors to guarantee animal movements (6–9). Accordingly, integration of neural circuits and body mechanics, referred to as neuromechanics, is one of the essential approaches to reveal motor control mechanisms (10–15).

Soft-bodied animals locomote by deforming their stretchable bodies (16). Compared to animals with hard skeletons, soft-bodied animals possess high flexibility and can move adaptively in complicated environments (17). This flexibility of soft-bodied animals suggests that output from neural circuits is not the sole determinant for locomotion. Animal’s physical properties such as stiffness and viscosity should also be involved in the dynamics of locomotion (9, 15) and have been measured experimentally (18–23). However, it is still challenging to reproduce locomotion in soft-bodied animals by mathematical models based on physical measurements. To tackle this issue, we used *Drosophila melanogaster* (*Drosophila*, hereafter) larvae as a model and attempted to build a neuromechanical model describing their crawling behaviour.

Forward crawling is the most predominant mode in fly larval locomotion (24, 25). By propagating segmental contraction from the posterior to anterior segments, larvae move forward (26–28), and neural circuits for crawling have been intensively examined (29–37). Previous simulation studies have succeeded in building models describing the propagative nature of crawling behaviour qualitatively (10,38–40). In contrast, the physical properties of larvae remained to be studied, and quantitative reproduction of crawling kinematics has not been achieved yet.

Here, we present a neuromechanical model to describe fly larval crawling behaviour based on their locomotion kinematics and soft-bodied biomechanics. First, we recorded forward crawling in the third-instar larvae whose segmental boundaries were labelled by a fluorescent protein and extracted kinematics parameters of crawling. Then using a tensile tester combined with optogenetics, we measured larval contraction force. Furthermore, larval viscoelasticity was measured by the tensile tester. The measurement indicated that the properties of the larval body were described better by the standard linear solid (SLS) model than the Kelvin-Voigt model previously used (10, 41). By incorporating these physical parameters with a neural circuit model built previously (10), we established a neuromechanical model for larval locomotion. By optimizing the neural circuit model parameters, we succeeded in building a neuromechanical model that reproduced the kinematics parameter extracted from larva measurements. This model could also reproduce optogenetic studies reported previously. In addition, crawling in a low friction condition was simulated, and its prediction was confirmed experimentally by analyzing floating larvae. Perturbation analyses in our model predicted the importance of intersegmental connections in speed control, contrasting to previous studies (10, 40). Based on our simulation results, we predicted the intersegmental connectivity in sister species in the genus *Drosophila*. Our model for larval crawling based on physical measurements provides a new approach to locomotion research in soft-bodied animals and soft robot engineering.

## Results

### Measurement of forward crawling behaviour in third-instar larvae

To reveal the physical mechanisms behind *Drosophila* larval crawling, we examined the kinematics of the behaviour and physical properties of the body. We used third-instar larvae because their size (3.53 ± 0.12 mm, n = 9 larvae) was large enough to measure physical properties as a soft material. While previous studies examined first- or second- instar *Drosophila* larvae and characterized the segmental kinematics (26,27,37,42,43), crawling behaviour in the third-instar larvae had not been investigated at a segmental scale but instead at a scale of the entire body (24,25,28,32,44). Thus, we examined the segmental dynamics in freely moving third-instar larvae.

First, we recorded the position of the segment boundaries in freely crawling larvae. To reliably trace the segmental boundaries of larvae, we expressed a GFP-tagged coagulation protein Fondue, which accumulates at the muscle attachment sites (45). We acquired time-lapse fluorescence images from the dorsal side of the larvae where longitudinal muscles span single segments (Figure 1A). From these fluorescence images, the segmental boundaries of the thoracic (T2-T3) and abdominal (A1-A8) segments were annotated (Figure 1B. See Materials and Methods for details.) We examined the displacement of each segment boundary during forward crawling. As previously reported in the first- and second-instar larvae cases, sequential displacement of the segmental boundary from the posterior to anterior segments was observed in third-instar larvae (Figure 1C). Four kinematic parameters were measured from the recording: Stride length, stride duration, intersegmental delay, and speed (Figure 1D). We obtained that stride length was 0.70 ± 0.05 mm, stride duration was 1.07 ± 0.12 sec, intersegmental delay was 0.09 ± 0.01 sec, and speed was 0.64 ± 0.02 mm/sec (n = 9 larvae).

**Figure 1.**
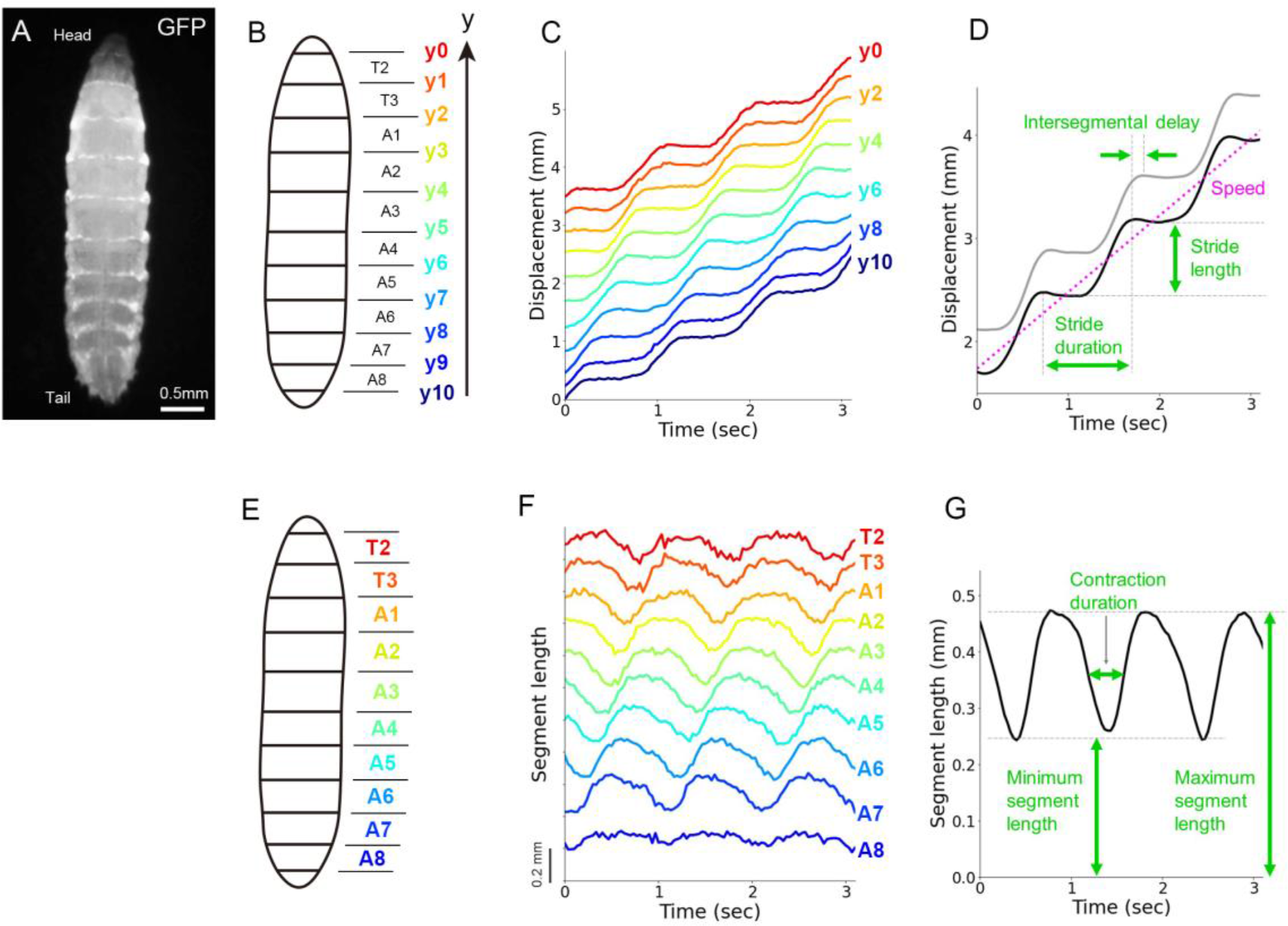
Characterization of forward crawling in third-instar larvae. (A) A fluorescence image of a third-instar larva with the anterior to the top. The segmental boundaries were visualized using *tubP-Gal4, UAS-fondue::GFP*. (B) A schematic of the segment boundaries (y0-y10) and the segment names (T2-A8). (C) Displacement of the segment boundaries during larval crawling. (D) Kinematics parameters based on segmental boundary dynamics. (E) A schematic of the segment names. (F) Segment length changes during larval crawling. (G) Kinematics parameters in segmental length change.

Next, we quantified the dynamics of segmental length change. Plots of the segmental length showed the propagation of segmental contraction in crawling larvae (Figure 1E and 1F). We measured three kinematic parameters (maximum segment length, minimum segment length, and contraction duration described in Figure 1G) and obtained that maximum segmental length was 0.41 ± 0.03 mm, minimum segmental length was 0.21 ± 0.02 mm, and contraction duration was 0.44 ± 0.03 sec (n = 9 larvae). To sum, we measured the seven kinematics quantities from segmental dynamics in freely crawling GFP-labelled third-instar larvae.

### Consistent kinematic properties among segments in larval crawling

To establish a mathematical model for larval crawling, we examined whether we could assume that all the segments possess similar kinematic properties or not. To this aim, we compared the dynamics of segmental length change (defined in Figure 1G) among all the segments (Figure 2A-2C). We noticed that the most posterior A8 segment had a shorter maximum length and smaller contraction range than the other segments (Figure 2B). This observation reflected the fact that the A8 segment was a specialized structure and different from the others in the surface area of the body wall and the number of muscles (46). In contrast, in the other segments, both the minimum and maximum lengths were comparable over the segments. (Figure 2B). Furthermore, contraction duration was also similar among the segments from T2 to A7 (Figure 2C). Accordingly, most segments (T2-A7) exhibited similar kinematic dynamics in contraction during crawling, which allowed us to model the larva as a chain of segments with the same kinematic properties.

**Figure 2.**
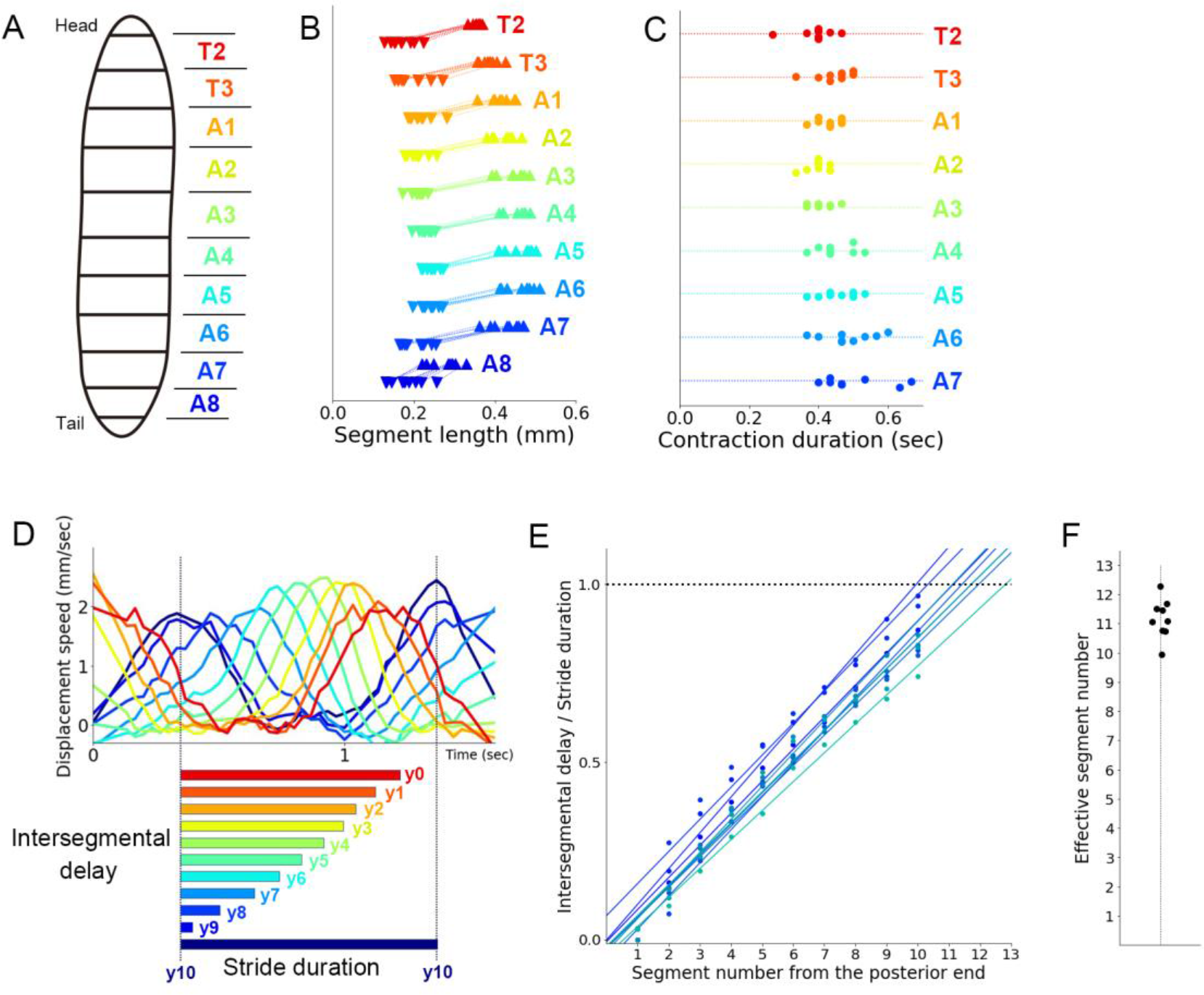
Consistent kinematics properties among segments in larval crawling. (A) A schematic of the segment names. (B) The range of segment length changes during larval crawling. Upward and downward triangles show the maximum and minimum length of the segments, respectively. (C) Contraction duration in larval segmental dynamics. (D) Displacement speed of larval segmental boundaries and schematics of their intersegmental delay. (E) Linear regression for larval intersegmental delays. (F) Effective segment numbers estimated by linear regression in (E).

Having confirmed the similarity among the body segments, we inferred the effective number of segments. First, we adapted a segment model based on the “piston phase” of larval peristalsis, where the motion of the head and tail were coupled (10, 26). When a peristaltic wave reached the head, the tail was forced to move forward, which induced the contraction of the posterior segment A8 for the next stride of crawling. Then, we estimated the effective number of segments by assuming that the displacement of the virtual anterior end was the same as that of the tail. To evaluate the propagation velocity, we obtained intersegmental delay by calculating the cross-correlation of displacements between the most posterior segment boundary (y10) and the other segments (y0-y9) (Figure 2D). The graph showed the propagation of segmental contraction from the posterior to the anterior segment. By plotting the intersegmental delay normalized by the stride duration, we noticed that the segmental contraction propagates at about a constant speed (Figure 2E). This observation allowed us to model the larval crawling as the propagation of segmental contraction at a uniform speed. Based on this assumption, we inferred the effective number of segments of the whole body where the head and tail were coupled. To this aim, we fitted the intersegmental delay data by linear regression. The intersection points between the fitting lines and the line that the normalized intersegmental delay equalled one corresponded to the event when the propagation of segmental contraction reached the anterior end and the subsequent propagation was initiated. From this calculation, we obtained 11.2 ± 0.2 as the effective number of segments (Figure 2F). Accordingly, the results indicated that the crawling larva could be modelled by a chain of 11 identical segments (Supplementary Figure 1).

### Measurement of larval contraction force

To establish a quantitative physical model for larval crawling, we attempted to measure contraction force and soft material properties of larvae which enabled us to derive a Newtonian description of crawling by linking force to motion. We measured contraction force with the tensile tester (Figure 3A). The tensile tester allowed us to measure and control both strain and stress of a sample material. We hooked an intact larva with insect pins and loaded it to the tensile tester. We measured the spontaneous contraction force in larvae and obtained values ranging from 1.4 mN to 2.7 mN (n=11 spontaneous contraction events by three larvae, Figure 3B and 3E). Although these values reflected the range of spontaneous contraction force, the larvae could not exhibit crawling locomotion during the measurement since their head and tail were hooked. Accordingly, we instead attempted to measure the maximum force that larvae could generate for crawling. To this aim, we adapted optogenetics to activate motor neurons reliably (Figure 3C). We expressed a light-sensitive cation channel protein Channelrhodopsin2 (47) in all of the larval type-I motor neurons (48) and illuminated each larva with blue light (455 nm, 5.7 nW/mm2) to induce the contraction of all the body wall muscles. Since we needed muscular force evoked by the CNS, we expressed Channelrhodopsin2 in motor neurons instead of muscles. The contraction force measurement with the optogenetic activation showed that the force ranged from 1.6 mN to 6.7 mN (n=44 optogenetic stimulations applied to seven larvae, Figure 3D and 3E). The variation in the measurement would be caused by differences in some uncontrollable conditions, including hooking states at the head and tail. Consequently, we adapted the maximum measured force of 6.7 mN as the maximum force (*F*_*Mmax*_) used in our neuromechanical model.

**Figure 3.**
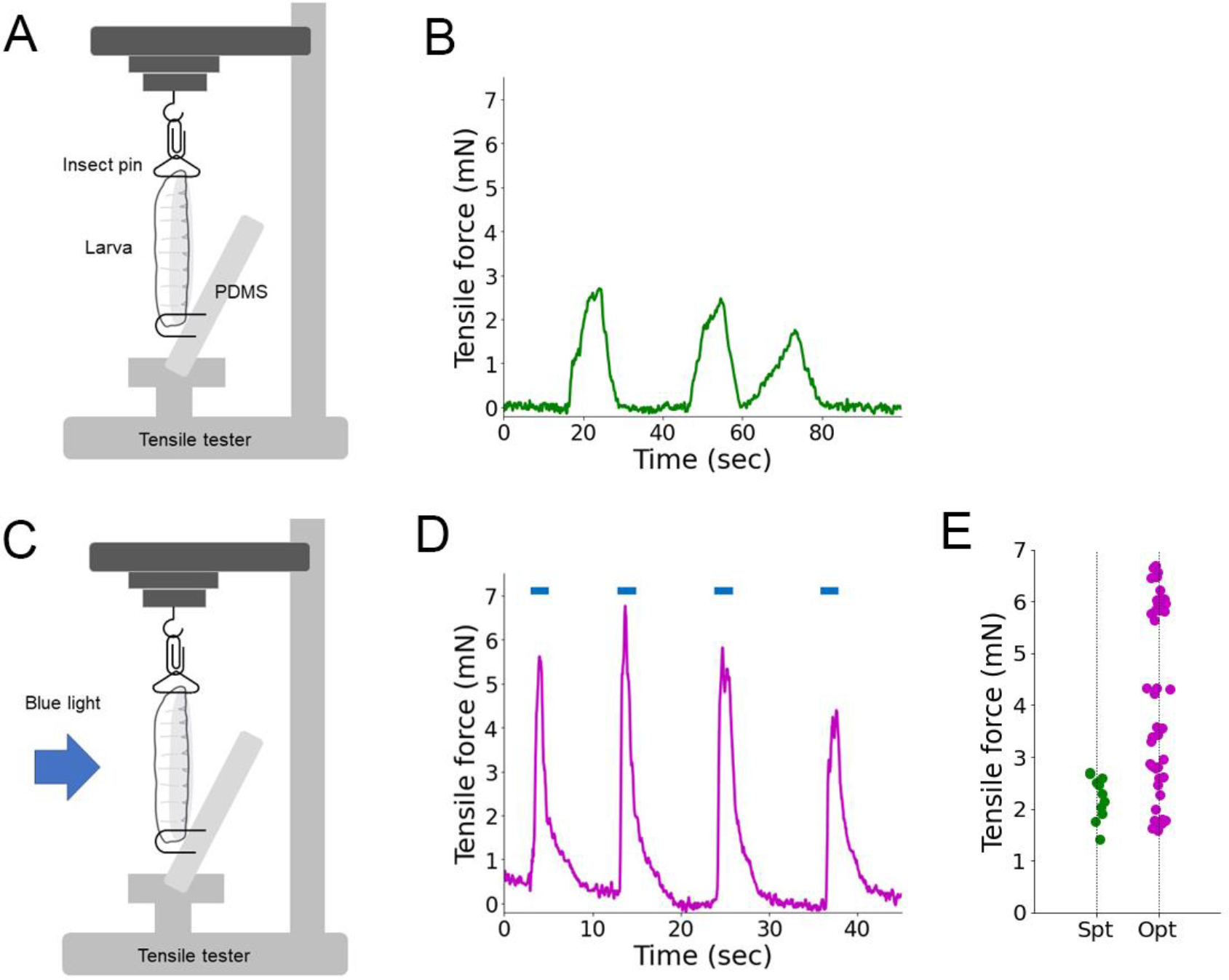
Measurement of contraction force in larvae. (A) A larva was hooked at the head and tail. The extension and tensile force of the larva were measured by a tensile tester. (B) A sample trace of spontaneous contraction force of a larva. (C) Optogenetic stimulation to a larva during measurement with the tensile tester. (D) A sample trace of larval force with optogenetic activation (shown in blue bars above the trace). (E) The tensile force of larvae generated by spontaneous contraction (“Spt”; n=11 contraction events from three larvae) and optogenetic activation (“Opt”; n=44 activations from seven larvae).

### Measurement of elastic and damping properties of larvae

Next, we analysed passive properties of the larval body, elasticity and viscosity, with the tensile tester (Figure 4A). The stretchable body wall and hydrostatic skeleton of fruit fly larvae were typical components in soft-bodied organisms (49). Previous studies reported material properties of soft-bodied animals, including the caterpillar *Manduca sexta* (18) and the earthworm *Lumbricus terrestris* (50, 51). Although previous theoretical analyses of fly larval locomotion assumed that each segment was equivalent to a pair of a spring (a linear elastic component) and a damper (a linear viscous component) (10,39,41), mechanical properties of *Drosophila* larvae were not measured experimentally. Accordingly, we measured the viscoelastic properties of fly larvae by methods used in material science.

**Figure 4.**
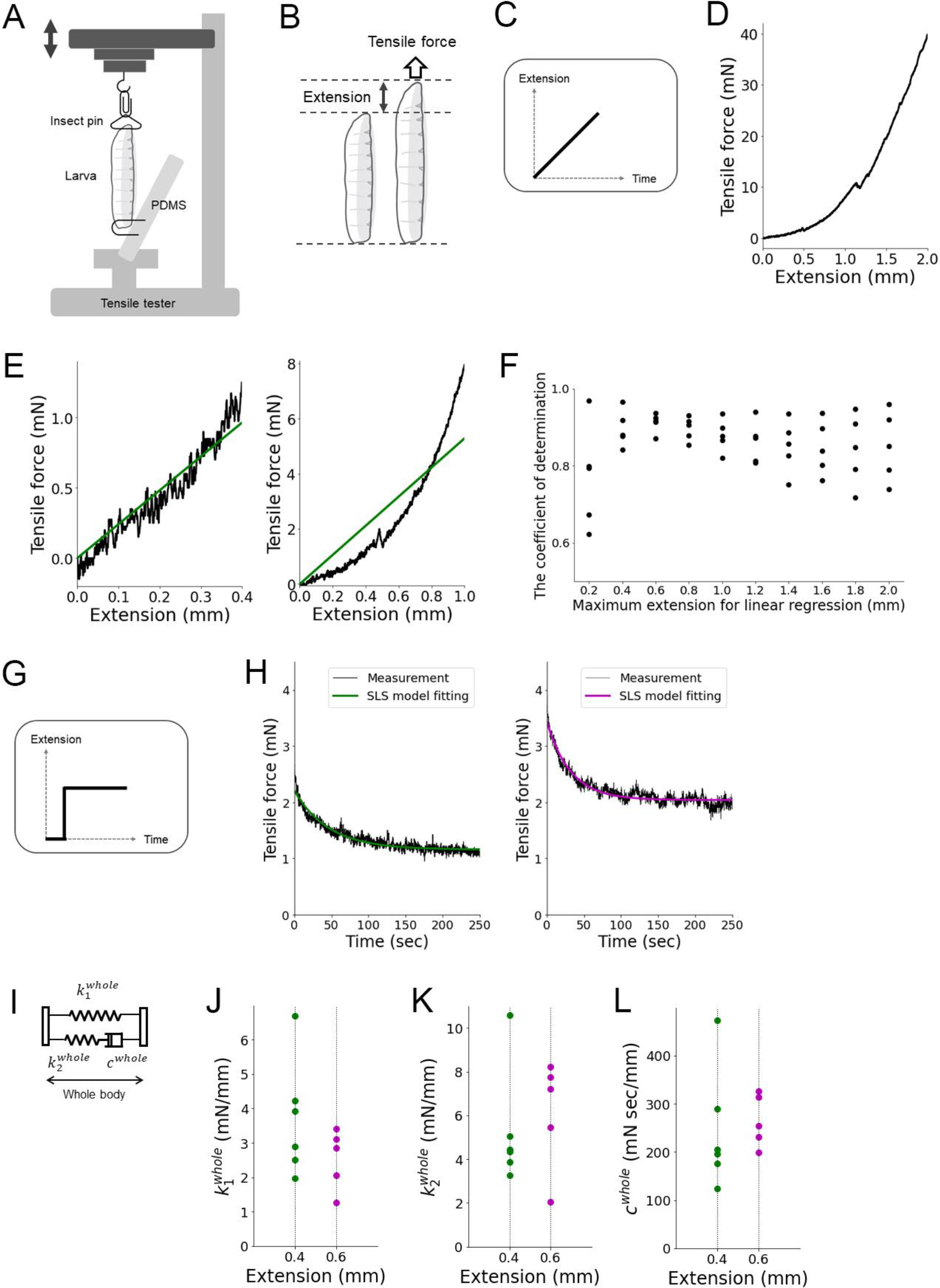
Measurement of viscoelasticity of the larval body. (A) Measurement and control of extension and tensile force of larval body with the tensile tester. The double-headed arrow indicates that the extension is changeable during the measurement. (B) A schematic of tensile force and extension of a larva. The left larva is not subjected to external forces while the right one is extended. The applied force to extend is the tensile force. (C) In the measurement of the relationship between extension and tensile force, the larvae were extended at a constant rate as shown in this panel, and the tensile force is being recorded. (D) An example trace of the relationship between extension and tensile force. (E) Plots of (D) in two different ranges of extension (Left: 0-0.4 mm, right: 0-1.0 mm). Green lines indicate linear regression lines in these ranges. (F) The coefficient of determination of linear regression in different ranges of extension. (G) In the stress-relaxation test, the larvae were extended quickly, and their length was kept constant. The tensile force under the constant extension was being recorded. (H) Example traces of stress-relaxation tests with the constant strain of 0.4 mm (left) and 0.6 mm (right). Fitting curves with the SLS model are shown in green (left) and magenta (right). (I) By the SLS model, the whole larval body can be described as two springs (spring constants: *k*_1_^*whole*^ and *k*_2_^*whole*^) and one damper (damping coefficient: *c*^*whole*^) (J-L) Scatter plots of *k*_1_^*whole*^, *k*_2_^*whole*^ (*K*), and *c*^*whole*^ (L) measured with the maximum strain of 0.4 mm (green) and 0.6 mm (magenta).

First, we analysed the range of extension within which we could assume linearity between the extension and tensile force of the larval body (Figure 4B). To analyse the relationship between the extension and tensile force of larvae, we stretched larvae at a constant speed (Figure 4C) and plotted tensile force against the extension (Figure 4D). To test the linearity, we used linear regression. Depending on the range of extension, goodness of fitting varies: Linear regression fitted better in the range from 0 mm to 0.4 mm extension (Figure 4E left) than in a longer range from 0 mm to 1.0 mm (Figure 4E right). To find a suitable extension range with good linearity, we calculated the coefficient of determinants, which was an indicator of predictability of the data by linear regression (Figure 4F). The plot showed that extension up to 0.4 mm or 0.6 mm exhibits high coefficients. Based on this observation, we concluded that we could assume linearity in the stretch ranging from 0 mm to 0.6 mm.

Next, we measured the viscoelasticity of larvae. Viscoelastic properties of soft materials could be acquired by measuring the relationship between stress (force loaded in the material) and strain (deformation of the material) (52). To obtain the viscoelastic properties of larvae, we conducted the stress relaxation test, one of the standard dynamic mechanical tests (52). In this test, a soft material sample was quickly extended to a certain length, and then the stress was being recorded over time (Figure 4G). Based on the linearity test above, we extended third instar larvae by 0.4 mm. The result of the stress relaxation test showed an exponential decay that was approaching a non-zero plateau at a later time (Figure 4H left). To extract physical properties from the stress relaxation data, a suitable physical model to describe the larval body was required. Among general mechanical structure models of linear combinations of springs and dampers, we found that the Maxwell model and the Kelvin-Voigt model, which were used previously to model larval musculature (10, 39), could not fit the experimental curve well since either exponential decay or residual elastic force was missing in these models (Supplementary Figure 2). In contrast, the standard linear solid (SLS) model, which combined the Maxwell model and a Hookean spring in parallel (52) (Figure 4I), fitted the experimental results well. Thus, we fitted this curve with the general SLS model (Figure 4H left). According to the SLS model, we obtained the two spring constants and one damping coefficient for the whole body of intact larvae by the extension of 0.4 mm (*k*_1_^*whole*^ = 3.7 ± 0.7 mN/mm, *k*_1_^*whole*^ = 5.3 ± 1.1 mN/mm, *c*^*whole*^ = (2.4 ± 0.5) × 10^2^ mN sec/mm, from n = 6 larvae, Figure 4J-4L). Consistent values were obtained by the extension of 0.6 mm (*k*_1_^*whole*^ = 2.5 ± 0.4 mN/mm, *k*_1_^*whole*^ = 6.1 ± 1.1 mN/mm, *c*^*whole*^ = (2.6 ± 0.2) × 10^2^ mN sec/mm, from n=5 larvae, Figure 4J-4L), which further supported the linear property of larvae. The two spring constants didn’t have a strong correlation (r = 0.49, p = 0.13, n = 11; Pearson’s Correlation Coefficient) suggesting that they could not be reduced to a single spring constant. To sum, we obtained the viscoelastic parameters of larvae, two spring constants and one damping coefficient, by the stress relaxation test. It should be noted that in larval crawling, each segment exhibited contraction instead of extension which we induced in the measurement. However, we could not measure the viscoelasticity of larvae during contraction since larvae were bent instead of compressed when pushed from both ends. Accordingly, we assumed larvae possess the same linear viscoelastic properties both in extension and compression. It should also be noted that in larval crawling each segment contracts by about 0.2 mm (Figure 2B) while in the stress-relaxation experiment each segment was extended by about 0.04 mm (0.4 mm / 11 segments). There could be some non-linearity in the viscoelastic properties of the larval body. However, in our model we adapted the viscoelasticity measured with 0.4 mm extension as the approximate values for our mathematical model of larval crawling.

### A neuromechanical model for larval crawling referring to the physical measurement

We established a mathematical neuromechanical model based on the physical measurements described above. We obtained seven kinematic values characterizing crawling locomotion (Figure 5A) from the measurement of fly larvae (Figure 1 and 2). Our aim was to build a model with a neural circuit and mechanical components to reproduce the observed results quantitatively. Regarding the mechanical part, based on the measurements, we estimated that the larva was a chain of eleven consistent segments (Figure 2), each of which was described by the SLS model (Figure 4 and Supplementary Figure 1). These assumptions indicated that the viscoelastic parameters for single segments in the model (two spring constants *k*_1_ and *k*_2_ and one damping coefficient *c*) should be 11 times larger than the ones for the whole body (*k*_1_ = 40.7 mN/mm, *k*_2_ = 58.3 mN/mm, and *c* = 2640 mN sec/mm).

**Figure 5.**
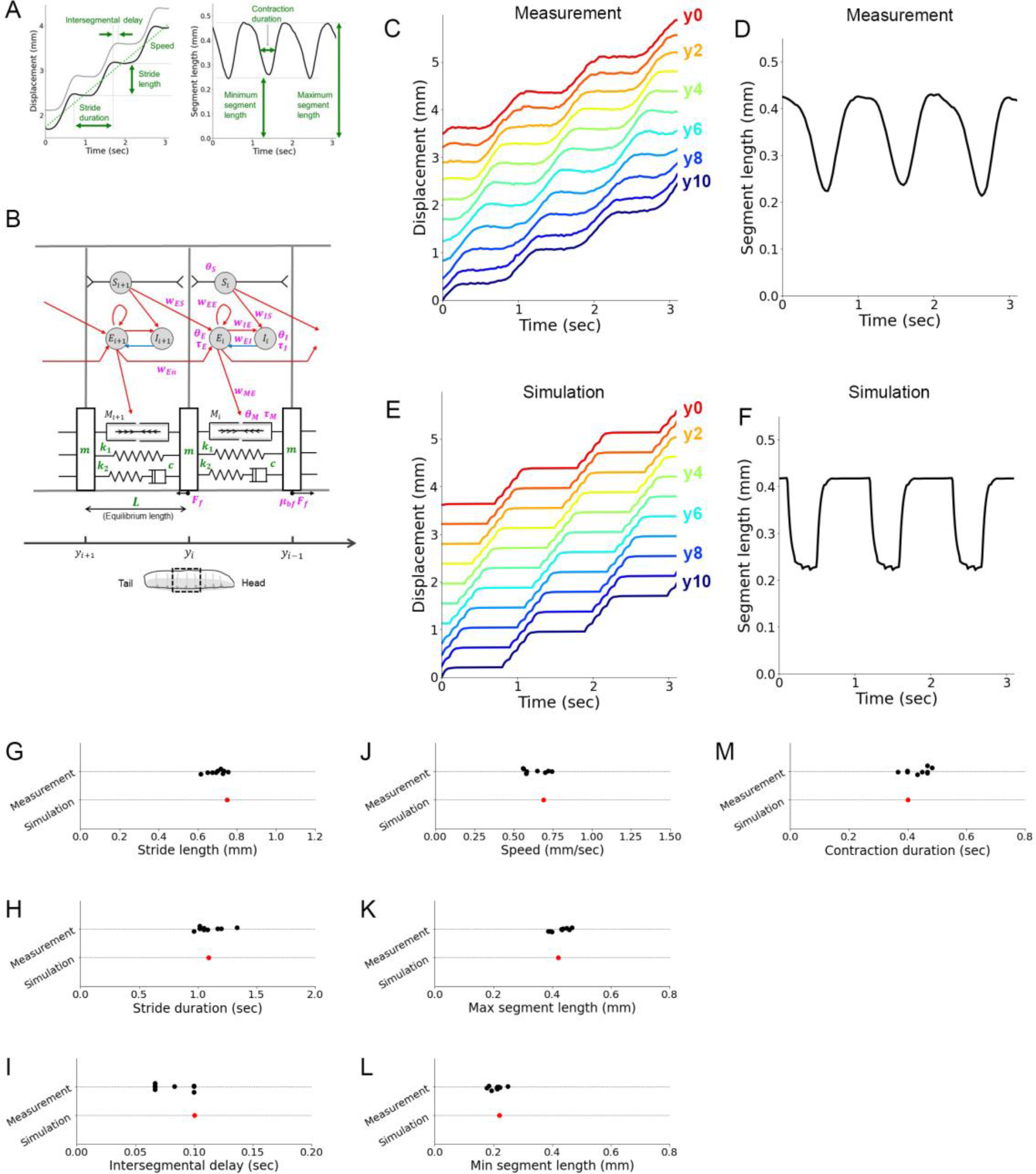
Reproduction of larval crawling by a neuromechanical model. (A) Kinematics parameters in segmental boundary dynamics (left, same as Figure 1D) and segmental length change (right, same as Figure 1G). (B) A schematic of our neuromechanical model for larval crawling. The neural circuit was referred to Pehlevan *et al*. (2016), although the parameter values in the model were different. Two segments in a larva (corresponding to the dotted box below) are drawn. Circles are populations of neurons: *E*_*i*_ is an excitatory neuron group, *I*_*i*_ is an inhibitory neuron group, and *S*_*i*_ is a sensory neuron group in the i-th segment. Excitatory and inhibitory connections are labelled by red and blue arrows, respectively. The body of the i-th segment is modelled by two springs (*k*_1_ and *k*_2_), one damper (*c*), and one muscle whose force is *M*_*i*_. Segment boundaries have masses (*m*) that feel friction with the substrate (*E*_*f*_ in forward motion and μ_*bf*_*F*_*f*_ in backward motion). The position of the i-th segment is denoted as *y*_*i*_ . (C) A kymograph of segmental boundaries during larval crawling (same as Figure 1C). (D) Change in segmental length during larval crawling (same data shown in Figure 1G). (E) A simulation result of kymograph of segmental boundaries. (F) Simulation result of segmental length. Segment labels and line colours in (C) and (E) correspond to those in Figure 1B and 1C. (G-L) Comparison of seven kinematics between parameters obtained from larval crawling (black dots, n = 9 larvae) and simulation (red dots). (G) Stride length. (H) Stride duration. (I) Intersegmental delay. (J) Crawling speed. (K) Maximum segment length. (L) Minimum segment length. (M) Contraction duration of single segments.

Furthermore, the force measurement indicated that the maximum muscular contraction force could be 6.7 mN (Figure 3). We modelled segmental boundaries as masses that were pulled and pushed by segmental muscles and dragged on the surface with friction (Figure 5B). We measured the mass of larvae (1.14 mg, the 99% confidence interval is [1.12, 1.15], n = 70 larvae) and calculated the mass of one segment by dividing the mass of the whole body by the number of segments to obtain 0.1 mg/segment. The length of larvae was 3.5 ± 0.1 mm (n = 9). We used all these values in our mechanical model (green parameters in Figure 5B). In contrast, as for the neural network part, we adapted a neural circuit model built by Pehlevan *et al*. (2016) based on Wilson-Cowan equations. This model was capable of generating propagation of waves from the posterior to anterior segments by interaction among excitatory and inhibitory neurons in the central nervous system (CNS), motor neurons, and sensory neurons (10). Although the parameters in this model were suggested, we incorporated the neural circuit framework and tuned its parameters to build a neuromechanical model based on physical measurement to reproduce larval crawling.

Aiming to reproduce the measurement results on larval crawling, we adjusted 15 parameters in the neural circuit model (magenta parameters in Figure 5B). The optimized set of the parameters (Supplementary Figure 3) could reproduce the measurement results of the kinematics (Figure 5C-5F). To assess this observation quantitatively, we compared the seven kinematics parameters (Figure 5A) between the measurement and simulation and found that all the parameters could be reproduced by the simulation to fit in the ranges of the measurement results (Figure 5G-5M). Accordingly, this neuromechanical model could quantitatively describe the segmental kinematics during forward larvae crawling.

The stress-relaxation results showed large variabilities on the order of 100% of the mean values (Figure 4J-4L). This variance could be caused by the uncertainty in the measurement or the variability in physical properties among larvae. Since we didn’t have independent measurement methods to obtain viscoelasticity, we couldn’t determine which one was the case. However, when we perturbed the spring constants by the order of 100% of the optimized value, crawling speed changed to be out of the measured range (Supplementary Figure 4). This observation suggested that the viscoelasticity would be consistent between larvae, and the variabilities in the stress-relaxation test would be caused by uncertainty in the measurement.

We adapted μ_*bf*_, the asymmetricity parameter between forward and backward friction as 10 considering the fact that a larva had denticle belts on the ventral side of the body, providing asymmetric friction with the ground. To test the dependency of crawling on this asymmetricity, we perturbed μ_*bf*_ (Supplementary Figure 5). The simulation results showed that crawling speed didn’t change even when μ_*bf*_ equalled one, meaning that the friction was symmetric between forward and backward. This observation indicated that the asymmetricity in the friction along the body axis had a small role in generating forward crawling.

As for the maximum muscular contraction force, we adapted the maximum force observed by optogenetic stimulation. To test whether we could use smaller muscular forces to reproduce larval crawling, we conducted a perturbation analysis on the maximum muscular force in our neuromechanical model (Supplementary Figure 6). The simulation results showed that crawling speed reduced as the muscular force decreased. Accordingly, this observation suggested that crawling speed was sensitive to the maximum muscular force and the measurement of contraction force was crucial to build a neuromechanical model for larval crawling.

We compared parameters in our model based on physical measurements with those in the previous simulation by Pehlevan *et al*. (2016). In Pehlevan *et al*. (2016), parameters were scaled by three values, the spring constant *k*, the equilibrium length of one segment *L*, and the relaxation time scale of the excitatory neurons τ_*E*_. Accordingly, we substituted these parameters with our measurement results (*k* = 50 mN/mm, *L* = 0.35 mm, and τ_*E*_ = 35 msec) and obtained the absolute values of the parameters. By comparison, we found that the neural couplings and relaxation times in the CNS were almost consistent between our results and the assumption in Pehlevan *et al*. (2016). On the other hands, physical properties showed clear contrast between them. The damping coefficient in our result (*c* ∼ 2640 mN sec/mm) was about 400 times larger than the previous assumption (10) (*c* ∼ 6 mN sec/mm). When we adapted the damping coefficient 400 times smaller than our value, crawling speed was affected to be doubled (crawling speed: 0.69 mm/sec in the optimized condition; 1.32 mm/sec when the damping coefficient c was 400 times smaller than the optimized value of *c*, Supplementary Figure 4C). And the maximum muscle contraction force in our results (*F*_*Mmax*_ = 6.7 mN) was 2.2 times smaller than the previous assumption (10) (*F*_*Mmax*_ = 15 mN). In addition, neural couplings between the CNS and either muscles or sensory neurons also exhibited differences: The coupling between the excitatory neurons and muscles in our model (*W*_*tE*_ = 540) was larger the previous assumption (*W*_*tE*_ = 1 in (10)), and the couplings between the sensory neurons and the CNS in our model (*W*_*Es*_ = *W*_*IS*_ 300) were also larger than the previous assumption (*W*_*Es*_ = *W*_*IS*_ = 1.95 in (10)). To sum, our mechanical model had larger viscosity and smaller muscular force whereas the circuit model had larger couplings between the CNS and their targets in the peripheral structures than the assumption in the previous theoretical work.

### The model reproduces previous observations in optogenetic experiments

To validate our model, we simulated two optogenetic perturbation studies reported previously and compared the results with the experimental observations (32, 53). First, by optogenetic silencing of motor neurons in a few segments, propagation of contraction from the posterior to anterior segments was arrested, and all the segments were relaxed (53). In the previous simulation study (10), although propagation was arrested by silencing the excitatory group in one segment, its neighbouring segment kept contracted instead of being relaxed. We silenced excitatory neurons in the A3 segment for 2 seconds in our neuromechanical model (Figure 6A-6F) and found that the silencing arrested the propagating wave (Figure 6E). Furthermore, all the segments were relaxed by this optogenetic silencing (Figure 6F), which is consistent with the experimental observation (53). Inada *et al*. (2011) demonstrated that crawling was resumed after removing optogenetic silencing (53). This phenomenon was not reproduced in our simulation: No wave was generated after optogenetic silencing (Figure 6E and 6F), although waves were resumed after a short period of silencing (Supplementary Figure 7). As pointed out in Pehlevan *et al*. (2016), this would be due to the lack of such a mechanism in the neural circuit model. To sum, our model successfully reproduced the observation that the silencing excitatory neurons, particularly motor neurons, in a single segment arrested the propagation of peristaltic waves and relaxed all the segments.

**Figure 6.**
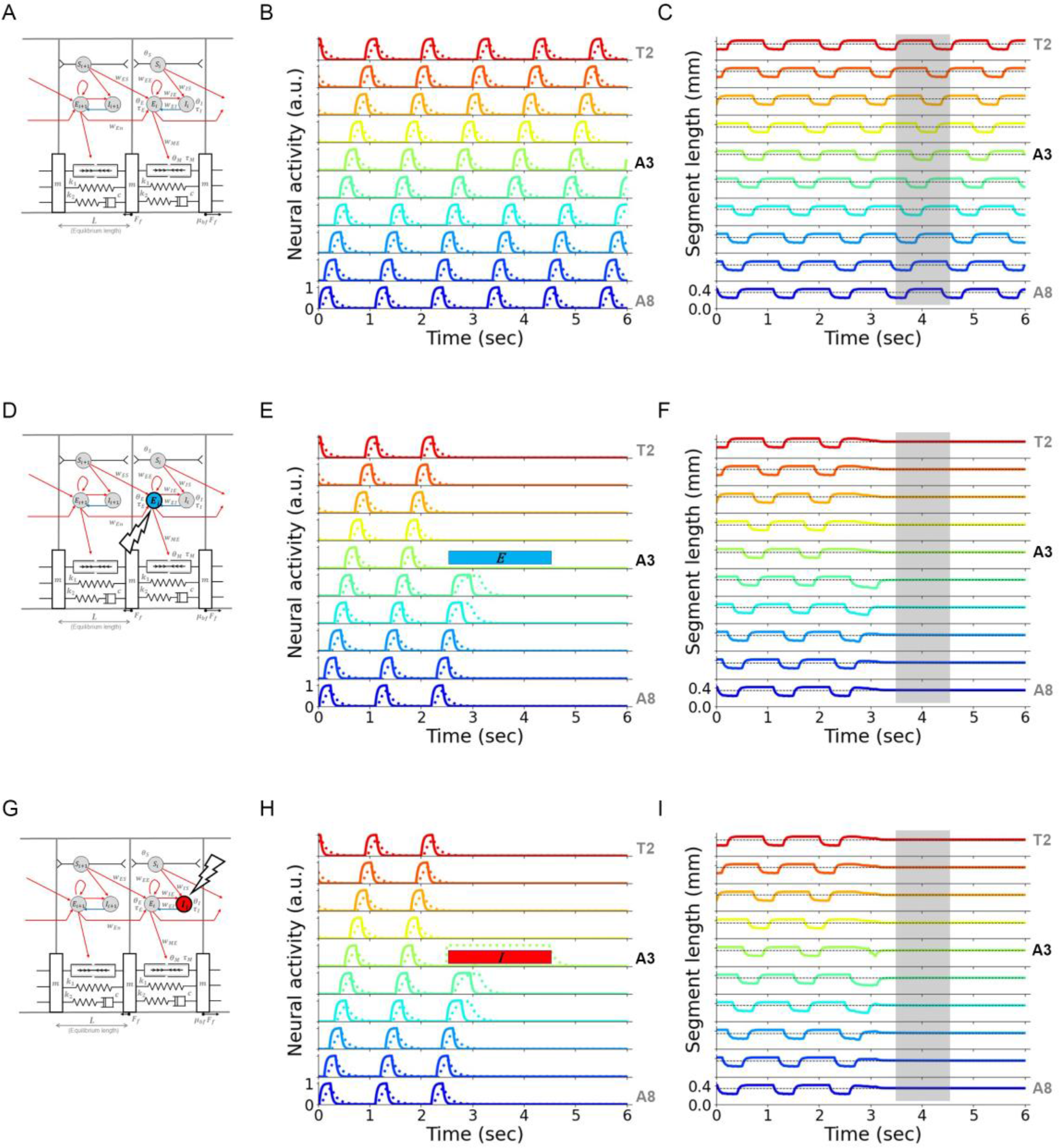
Reproduction of optogenetic experiments by the mathematical model. (A) A schematic of two segments in the mathematical models. (B) Activity of excitatory (solid lines) and inhibitory neurons (dotted lines) in A8-T2 segments by the model without optogenetic perturbation. (C) Traces of segment length in (B). Dotted lines denote the equilibrium length of the segments. A time period of 3.5-4.5 seconds is shaded to compare with (F) and (I). (D) A schematic of optogenetic silencing of excitatory neurons in a single segment shown by a blue disk. (E) Neural activity with the optogenetic silencing of excitatory neurons. Excitatory neurons in the A3 segment were silenced for 2 seconds, marked by a blue bar. Waves were arrested after the optogenetic silencing. (F) Traces of segment length in (E). Dotted lines denote the equilibrium length of the segments. In the later phase of the optogenetic silencing (the shaded region), the length of all the segments returned to the equilibrium length. (G) A schematic of optogenetic activation of inhibitory neurons in a single segment shown by a red disk. (H) Neural activity with the optogenetic activation of inhibitory neurons. Inhibitory neurons in the A3 segment were activated for 2 seconds, marked by a red bar. Waves were arrested after the optogenetic activation. (I) Traces of segment length in (H). Dotted lines denote the equilibrium length of the segments. In the later phase of the optogenetic activation of inhibitory neurons (the shaded region), the length of all the segments returned to the equilibrium length. Plot colours in B, C, E, F, H, and I correspond to those in Figure 1F.

Next, we tested the effect of activation of inhibitory neurons on crawling behaviour. A previous study reported that activation of inhibitory premotor interneurons PMSIs arrested wave propagation and caused the relaxation of the whole body (32). Therefore, we activated inhibitory neurons in the A3 segment in the neuromechanical model (Figure 6G). By this treatment, waves were arrested (Figure 6H), and the whole segments returned back to their equilibrium length (Figure 6I). Accordingly, our neuromechanical model successfully reproduced the two optogenetics experiment results.

### Measurement of forward crawling behaviour in a low-friction condition and validation of the model

Friction was a reaction force of a propelling force that drove larval movement. Although friction was necessary to move the centroid of larvae, it was unclear whether friction was required to generate the patterned segmental motion. In the previous theoretical study, two phenomena in segmental dynamics in a low friction condition were simulated: Each segment contracted more, and intersegmental propagation delay became longer (10). Our neuromechanical model tested the segmental dynamics in no friction condition (Figure 7A- 7H). The simulation results showed that the range of segment contraction didn’t change even in the no friction condition (Figure 7C – 7F). Also, the duration of segmental contraction was not affected either (Figure 7G). Furthermore, with respect to the wave propagation, intersegmental delay was not changed in no friction condition (Figure 7H). Accordingly, in our mathematical model, segmental dynamics was robust in the no friction condition.

**Figure 7.**
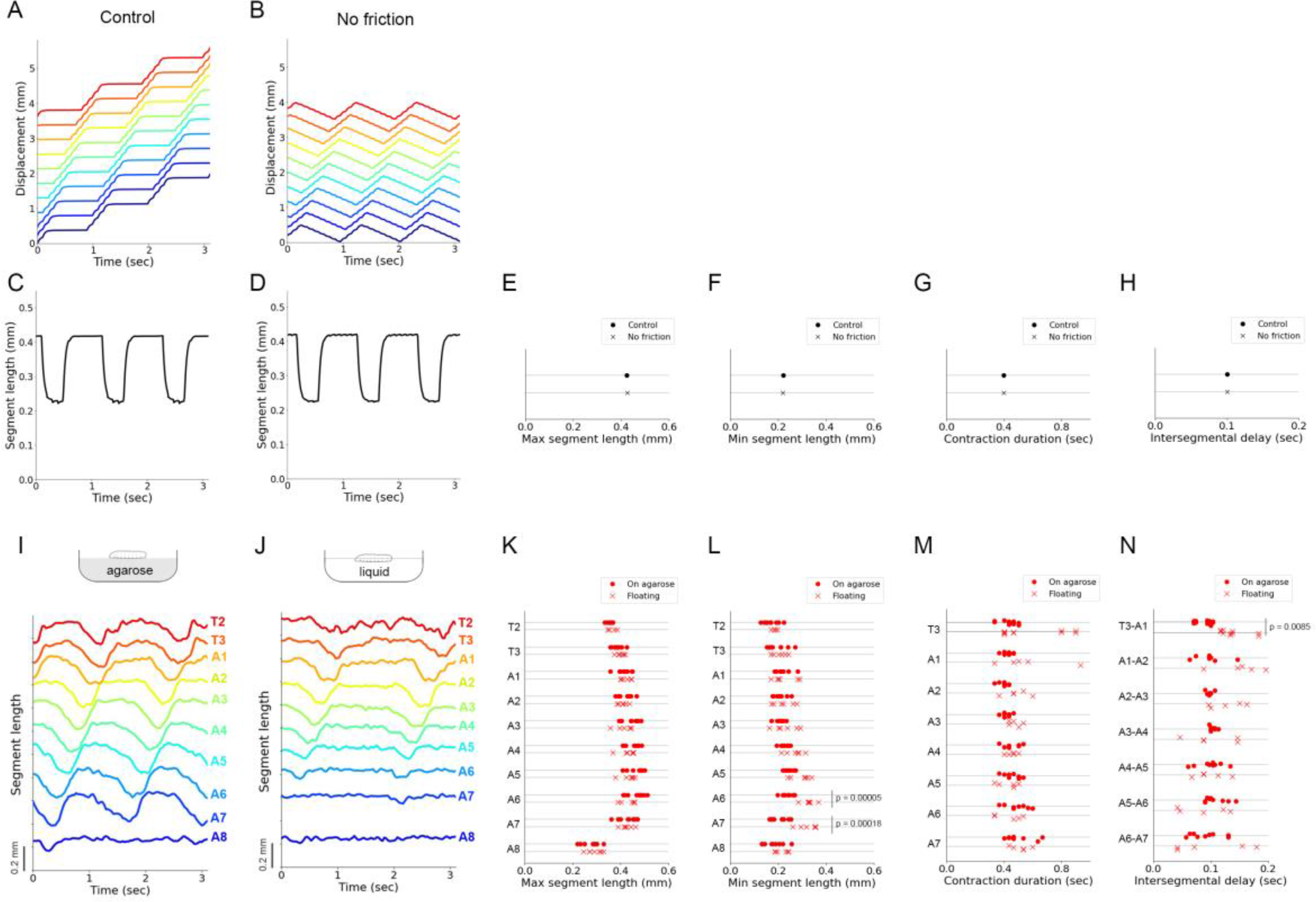
Mathematical simulation and experimental analysis of crawling behaviour in low friction conditions. (A and B) Simulation results of kymograph in the optimized condition (A) and the absence of friction (B). (C and D) Simulation results of segmental length in the optimized condition (C) and the absence of friction (D). (E-H) Comparison of four kinematic parameters in simulations between in the optimized condition (Control) and the absence of friction (No friction). (E) Maximum segment length. (F) Minimum segment length. (G) Contraction duration. (H) Intersegmental delay. (I and J) Plots of segment length measured by the experiments of larvae on agarose (I) and those floating in liquid (J). (K-N) Comparison of four kinematic parameters obtained by the experiments of larvae on agarose (On agarose) and larvae in a low friction condition (Floating). The p-values show the result of Student’s t-test. (K) Maximum segment length. (L) Minimum segment length. (M) Contraction duration. (N) Intersegmental delay.

To reveal the roles of friction on larval crawling, we measured body segment kinematics in crawling larvae in a low-friction environment. We used a high concentration sugar solution (66% w/w sucrose) and floated third-instar larvae on it. To visualize segment boundaries of larvae, we used *tubP-Gal4, UAS-fondue::GFP* fly larvae as described above (Figure 1A). We compared the crawling motion between on the agarose substrate (Figure 7I) and in the sugar solution (low friction environment) (Figure 7J). Floating larvae could exhibit crawling behaviour (Figure 7I), suggesting that the interaction with a substrate was not required to generate peristaltic motion. They could not move forward in the sugar solution probably due to lack of the propelling force that was the reaction force of friction or because of viscous resistance in the sugar solution. The number of waves was fewer in floating larvae than larvae on an agarose substrate (agarose: 45.5 ± 3.5 crawls/min, n = 6 larvae; floating: 6.8 ± 2.6 crawls/min, n = 6 larvae; p = 9.0 × 10^−6^, Welch’s t-test; Supplementary Figure 8A).

This observation implied that friction could be involved in the initiation of crawling. This phenomenon could not be observed in our neuromechanical model (Supplementary Figure 8B) probably because our model didn’t include a mechanism to control the initiation of waves independent of the completion of waves. How each crawling wave was initiated and how the temporal gap between consecutive waves was regulated remained important open questions. Since our focus in this study was on the segmental and intersegmental dynamics, we analyzed the kinematics of every single peristaltic wave.

First, we examined the segmental dynamics in the low friction condition and found that the range of segment contraction didn’t change in most of the segments in floating larvae (Figure 7K and 7L). In addition, the contraction duration of each segment was not affected in the floating condition (Figure 7M). These observations were consistent with our model results (Figure 7E-7G). In the posterior segment A6 and A7, however, there was a difference between larvae on agarose and floating larvae. The range of contraction at the posterior segments was reduced in floating larvae (Figure 7L). Since forward crawling started at the posterior segments, this observation implied that friction would be crucial to initiate crawling. This implication was consistent with the observation of the reduction in crawling numbers in floating larvae (Supplementary Figure 8A). Second, we analyzed the intersegmental delay in the floating larvae. In most segments, intersegmental delay was consistent between larvae on agarose and floating larvae (Figure 7N). However, the intersegmental delay between T3 and A1 was increased in floating larvae. Since this phenomenon was observed in the limited anterior segment, it would be possible that other behaviour, including head-sweeping that occurs in the anterior segments, was affected in the low friction condition. To sum, the kinematics in larval crawling was almost similar between on the agarose substrate and in low-friction condition. This observation was consistent with the result of our neuromechanical model.

### Contribution of sensory feedback and CNS

The CNS and proprioception were both involved in locomotion. The significant contribution of sensory feedback in larval crawling was shown experimentally (Hughes and Thomas 2007). This observation raised a hypothesis that proprioception could cause the propagation signal to the neighbouring segment through intersegmental sensory feedback (44). A simulation based on this assumption showed that the model circuit with intersegmental feedback could normally generate propagation waves without an intersegmental connection in the CNS (10). On the other hand, several key interneurons regulating larval locomotion had been identified (32,54–56), suggesting that interneurons in the CNS should be involved in wave propagation. Aiming to reproduce the experimental observation of the involvement of the CNS in crawling, we tested the contribution of intersegmental sensory feedback and the intersegmental connection in the CNS to peristaltic motion by our neuromechanical model (Figure 8). When the sensory feedback was silenced (*W*_*ES*_ = 0), crawling speed was reduced (Figure 8B, 8D, and 8H). This was consistent with previous simulation result (10) and the experimental result (44). When the intersegmental central connection was blocked (*W*_*En*_ = 0), the speed of crawling was decreased too (Figure 8B, 8F, and 8I). This result was also consistent with observations that interneurons in the CNS were involved in crawling speed. While the speed was reduced in both cases, the neural mechanisms would be different. When sensory feedback was blocked, the activity of inhibitory neurons was attenuated (Figure 8C and 8E). Meanwhile, when the intersegmental central connection was blocked, the activity of inhibitory neurons was enhanced (Figure 8C and 8G). This observation suggested that sensory feedback and the CNS were involved in the speed control by distinct mechanisms. To examine their roles in detail, we perturbed *W*_*ES*_ and *W*_*En*_ from the optimal values. The speed of crawling was robust to the change of *W*_*ES*_ (Figure 8H), whereas increase in *W*_*En*_ led to drastic change in crawling speed (Figure 8I). In particular, crawling speed depended on *W*_*En*_ nonlinearly with a peak. To sum, this simulation result was consistent with the experimental observations that interneurons in the CNS should be involved in the regulation of crawling behaviour.

**Figure 8.**
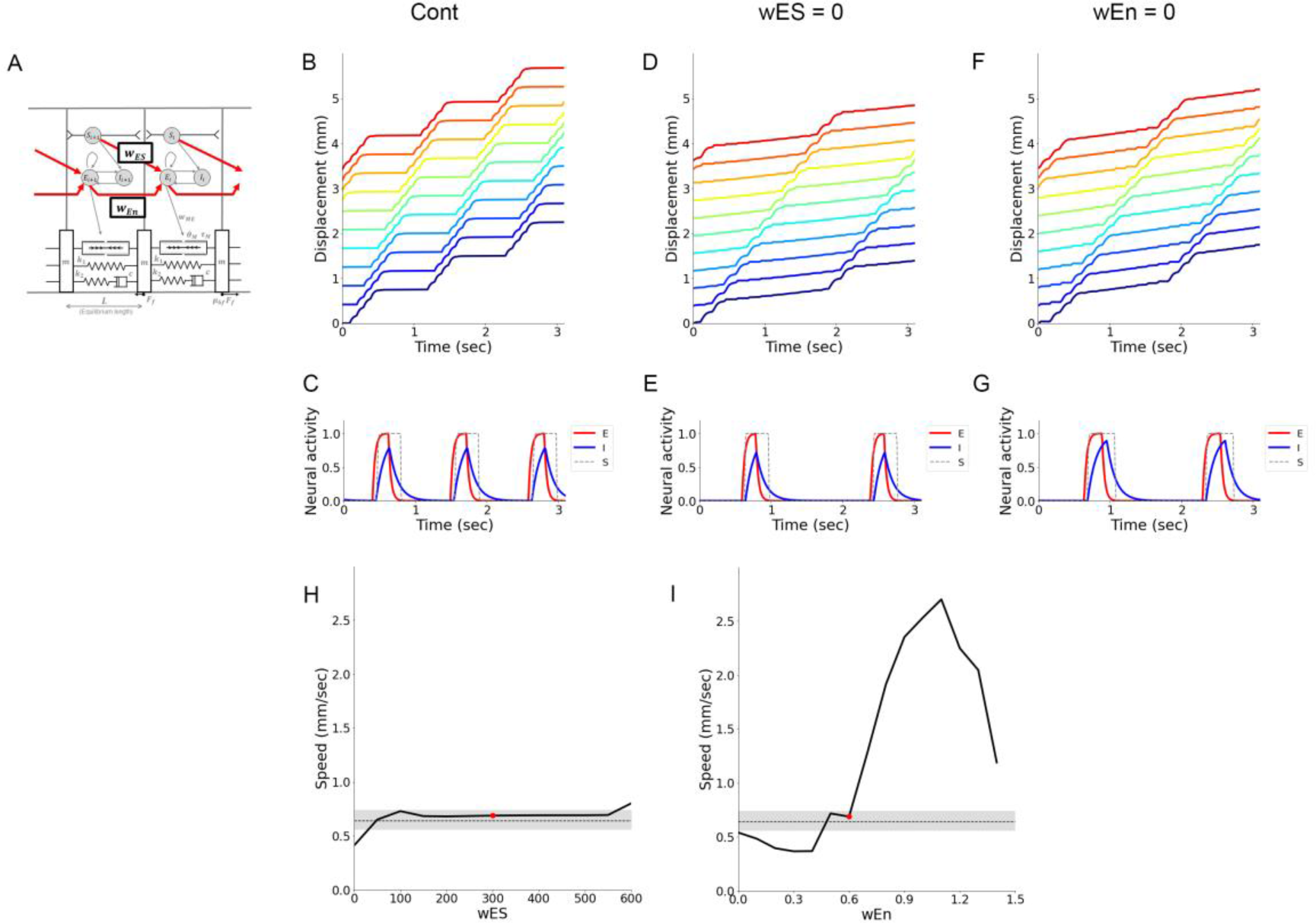
Sensory feedback and central connectivity are both involved in controlling crawling speed. The weights of intersegmental proprioceptive sensory feedback *W*_*ES*_ and intersegmental coupling *W*_*En*_ in the neuromechanical model. (B and C) Simulation in the optimized condition. Segmental boundary kymograph. (C) Activity of excitatory (red line), inhibitory (blue line), and sensory neurons (dotted lines) in a single segment. (D and E) Simulation in the absence of proprioceptive sensory feedback. (D) Segmental boundary kymograph. (E) Activity of excitatory (red line), inhibitory (blue line), and sensory neurons (dotted lines) in a single segment. (F and G) Simulation in the absence of intersegmental connections. (F) Segmental boundary kymograph. (G) Activity of excitatory (red line), inhibitory (blue line), and sensory neurons (dotted lines) in a single segment. (H and I) Plots of speed as proprioceptive sensory feedback *W*_*ES*_ (H) or intersegmental coupling *W*_*En*_ was perturbed (I). Grey shaded regions show the range of speed observed in the experiment with third-instar larvae. Red dots indicate the optimized simulation condition.

Next, we analyzed the interaction between intersegmental sensory feedback and intersegmental connection in the CNS. We perturbed *W*_*En*_ and *W*_*ES*_ simultaneously and plotted the displacement of larvae in a two-dimensional plot (Figure 9A). The plot showed that crawling could be realized in the wide area of this parameter space. When *W*_*En*_ was larger than the value in the optimized condition (0.6), *W*_*En*_ played a dominant role in speed control compared to *W*_*ES*_ (Figure 9B). On the other hand, in the regime of smaller *W*_*En*_, both *W*_*En*_ and *W*_*ES*_ were involved in crawling speed. This result indicated the strength of central connectivity in the CNS should play a pivotal role to control the speed of crawling behaviour.

**Figure 9.**
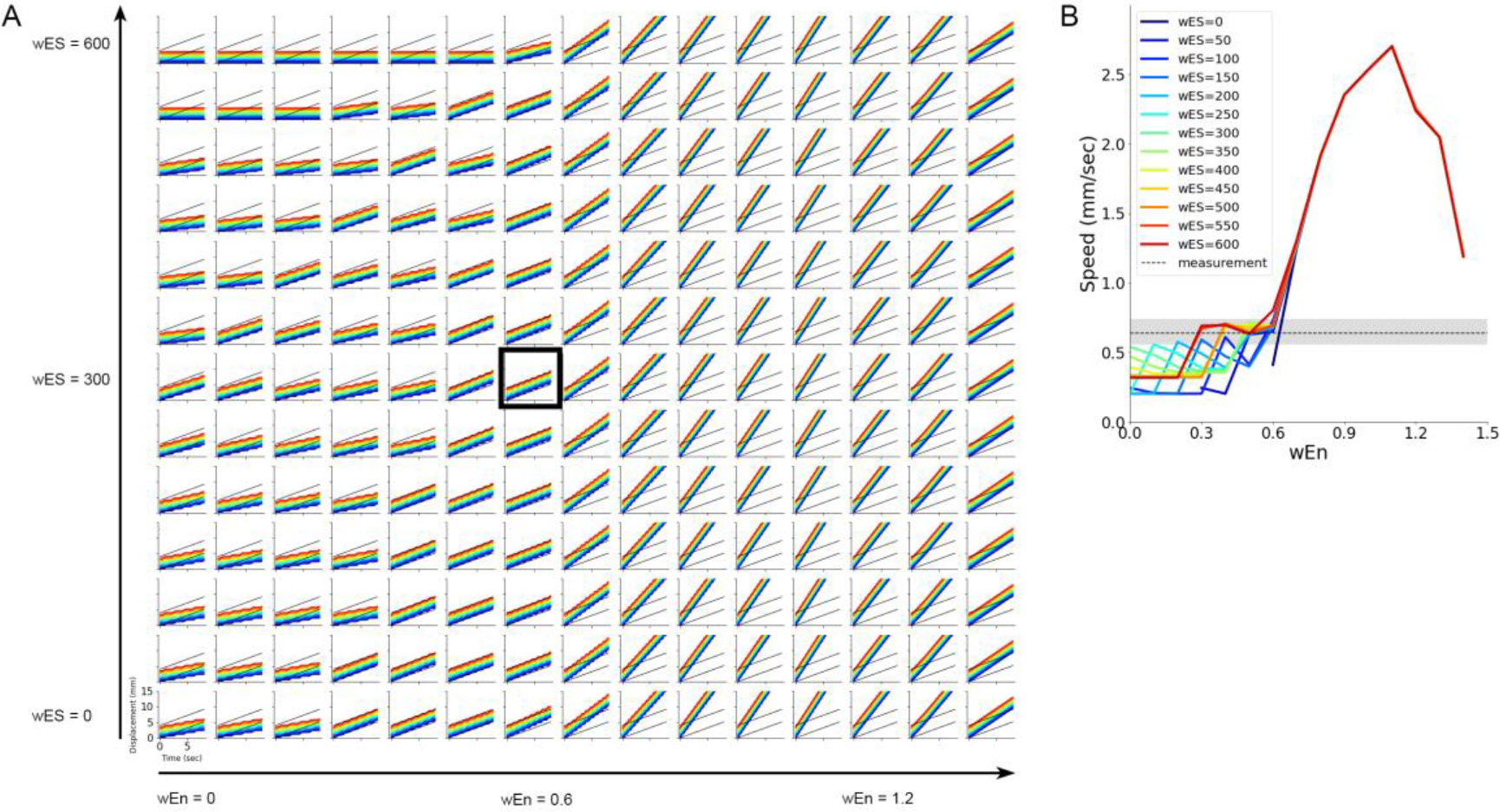
Interaction between *W*_*Es*_ and *W*_*En*_ in controlling crawling speed. (A) Plots of segment displacement in varied combinations of proprioceptive sensory feedback *W*_*ES*_ (along the vertical axis) and intersegmental coupling *W*_*En*_ (along the horizontal axis). The plot enclosed in a black rectangle corresponds to the optimized condition. (B) A plot of speed as both proprioceptive sensory feedback *W*_*ES*_ and intersegmental coupling *W*_*En*_ were perturbed. A grey line and a shaded region show the average and the range of speed observed in the experiment with third-instar larvae, respectively.

### Contribution of excitatory and inhibitory neurons in crawling

Based on the observation suggesting a significant contribution of the CNS to crawling behaviour, we next examined the involvement of excitatory and inhibitory neurons in larval locomotion. To this aim, we perturbed intrasegmental connection weight: *W*_*EE*_ for excitatory connections and *W*_*EI*_ for inhibitory connections (Figure 10A-10G). Crawl speed decreased when blocked excitatory connections (Figures 10B and 10D). Since the traces of excitatory and inhibitory neurons were different from those in blocking intersegmental excitatory connections (Figure 8E and 10E), the speed reduction mechanism should be different between silencing the intrasegmental excitatory connections and intersegmental excitatory connections. On the other hand, when the activity of inhibitory neurons was suppressed, larvae couldn’t exhibit crawling (Figure 10F). In this case, all the neurons exhibited tonic hyperactivity due to a lack of inhibition (Figure 10G). To further reveal the involvement of *W*_*EE*_ and *W*_*EI*_, we perturbed them in the range where larvae could exhibit crawling. The plots showed that speed was related to *W*_*EE*_ and *W*_*EI*_ in non-monotonical manners (Figure 10H and 10I). This observation indicated that both excitatory and inhibitory neurons controlled crawling speed.

**Fig. 10.**
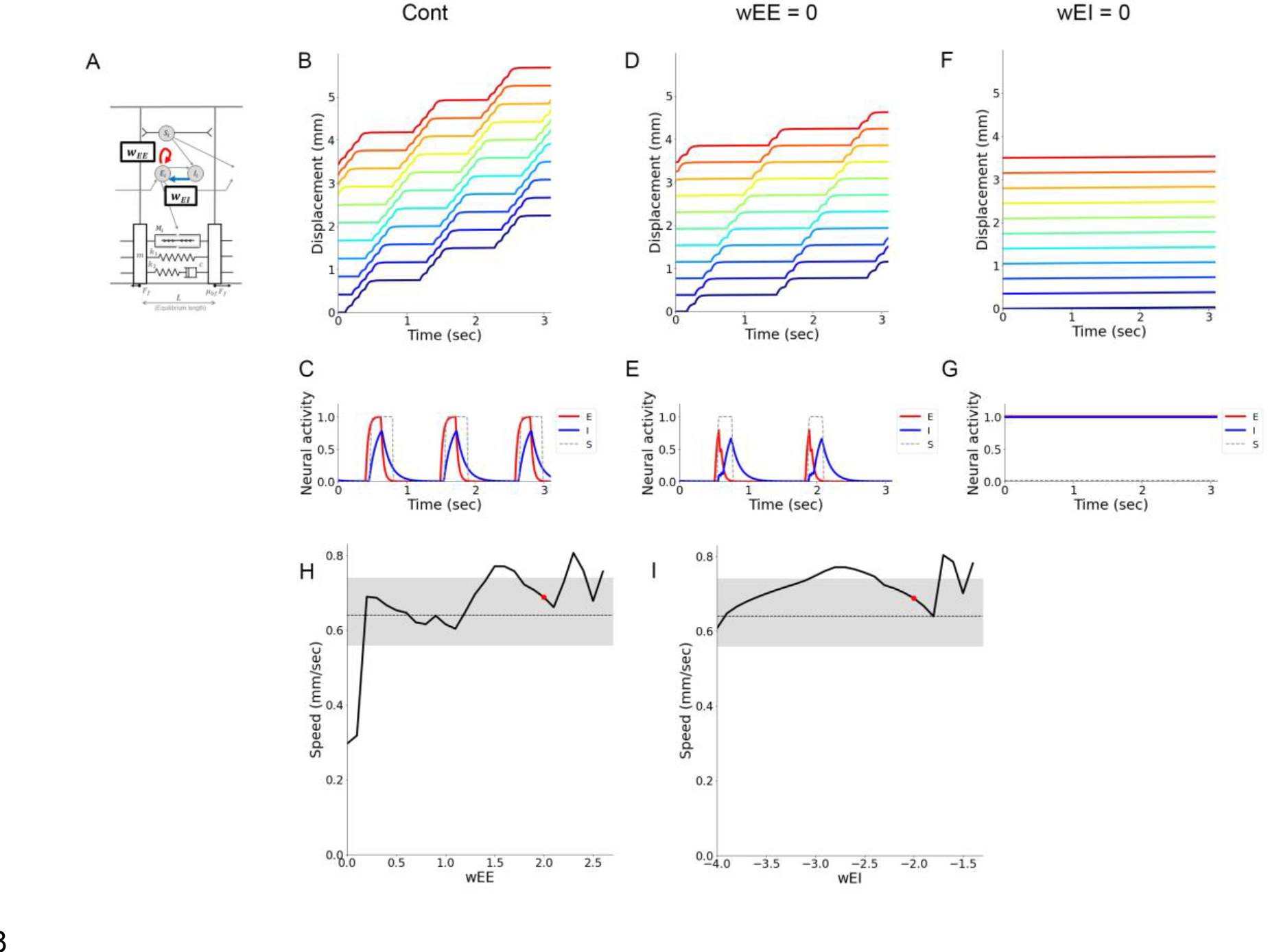
Intrasegmental excitatory and inhibitory connections are both involved in controlling crawling speed. (A) The weights of intrasegmental connections between excitatory neurons (*W*_*EE*_) and intrasegmental connections from inhibitory neurons to excitatory neurons (*W*_*EI*_) in the neuromechanical model. (B and C) Simulation in the optimized condition. (B) Segmental boundary kymograph. (C) Activity of excitatory (red line), inhibitory (blue line), and sensory neurons (dotted lines) in a single segment. (D and E) Simulation in the absence of intrasegmental connections between excitatory neurons. (D) Segmental boundary kymograph. (E) Activity of excitatory (red line), inhibitory (blue line), and sensory neurons (dotted lines) in a single segment. (F and G) Simulation in the absence of intrasegmental connections from inhibitory neurons to excitatory neurons. (F) Segmental boundary kymograph. (G) Activity of excitatory (red line), inhibitory (blue line), and sensory neurons (dotted lines) in a single segment. (H and I) Plots of speed as the weight of intrasegmental connections between excitatory neurons *W*_*EE*_ (H) or the weight of connections from inhibitory neurons to excitatory neurons *W*_*EI*_ was perturbed (I). Grey shaded regions show the range of speed observed in the experiment with third-instar larvae. Red dots indicate the optimized simulation condition.

Next, we analyzed the interaction between excitatory and inhibitory neurons. Displacement of larvae with different values of *W*_*EE*_ and *W*_*EI*_ was plotted two-dimensionally.

The diagram indicated that when *W*_*EE*_ was large and *W*_*EI*_ was close to zero (the upper right side of the diagram), larvae couldn’t generate crawling (Figure 11A). On the other hand, when *W*_*EE*_ was small (weak excitatory connection) or *W*_*EI*_ was small (strong inhibitory connection) that corresponded to the lower-left region of the diagram, crawling speed was reduced. Interestingly, as long as *W*_*EE*_ and *W*_*EI*_ were balanced (the diagonal region of the diagram), crawling speed was consistent. The deviation in speed when *W*_*EE*_ and *W*_*EI*_ were perturbed was within the range of variability observed experimentally (Figure 11B). These results indicated two points: The balance between excitatory and inhibitory connections within segments was critical to generate peristaltic motion. And, as long as they were balanced, crawling speed would not change by perturbation in intrasegmental connection weights.

**Fig. 11.**
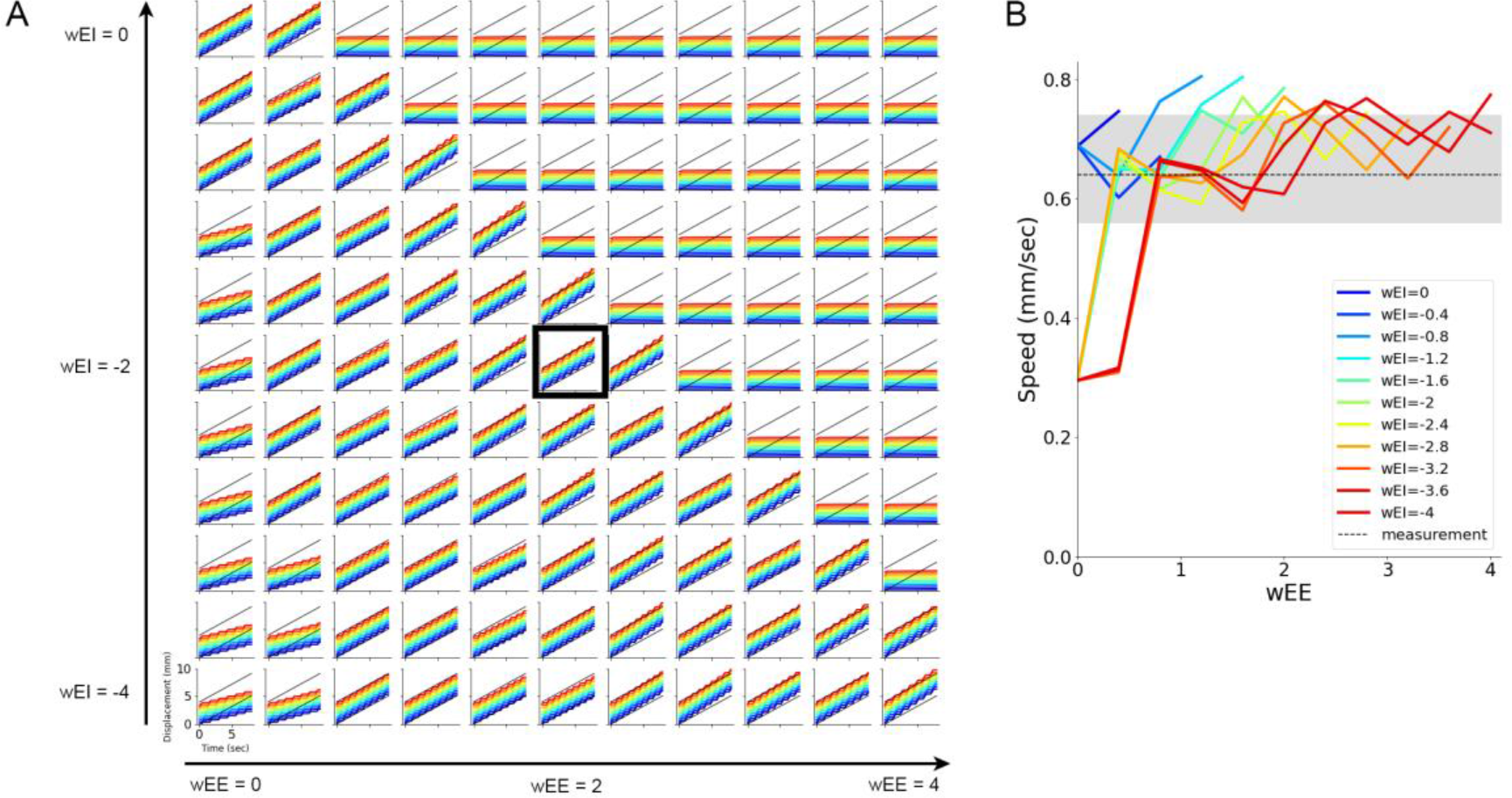
Interaction between *w*_*EE*_ and *w*_*EI*_ in controlling crawling speed. (A) Plots of segment displacement in varied combinations of the weights of intrasegmental connections between excitatory neurons *W*_*EE*_ (along the horizontal axis) and from inhibitory neurons to excitatory neurons *W*_*EI*_ (along the vertical axis). The plot enclosed in a black rectangle corresponds to the optimized condition. (B) A plot of speed as both *W*_*EE*_ and *W*_*EI*_ were perturbed. A grey line and a shaded region show the average and the range of speed observed in the experiment with third-instar larvae, respectively.

### Prediction in the neural network in sister species of *Drosophila melanogaster*

Comparison between the contribution of intersegmental (Figure 9B) and intrasegmental (Figure 11B) connections to crawling speed suggested that intersegmental connections in the CNS had dominant roles in regulating crawling speed. We noticed that this observation could allow us to predict neural circuits for fly larval crawling. A recent study reported an inter-specific variation in crawling speed among eleven species in the genus *Drosophila* (57). This study showed that some sister species crawled faster than *Drosophila melanogaster*, and the others locomoted slower. We attempted to make a prediction about the intersegmental connection in the CNS of these species based on their crawling speed. To predict the property of neural circuits in these sister species, we overlaid the speed data of the species on the graph showing speed dependency on *W*_*En*_ (Figure 12). To compensate the difference in experimental conditions between this study and Matsuo *et al.* (2021), the speed of each species was normalized based on the speed of *Drosophila melanogaster*. To show the dependency on *W*_*En*_, the dependency of *W*_*ES*_ was averaged out from Figure 9B. As a result, all of the eleven horizontal lines demonstrating the crawling speed of sister species crossed to the speed-*W*_*En*_ relation curve (Figure 12). This plot suggested that some species showing faster crawling, *Drosophila pseudoobscura* for example, should have strong intersegmental connections in the CNS while those crawling slower such as *Drosophila mojavensis,* should have weaker intersegmental central connections. This prediction would be testable by interspecific comparison based on the anatomy of their central nervous system or physiological recording.

**Fig. 12.**
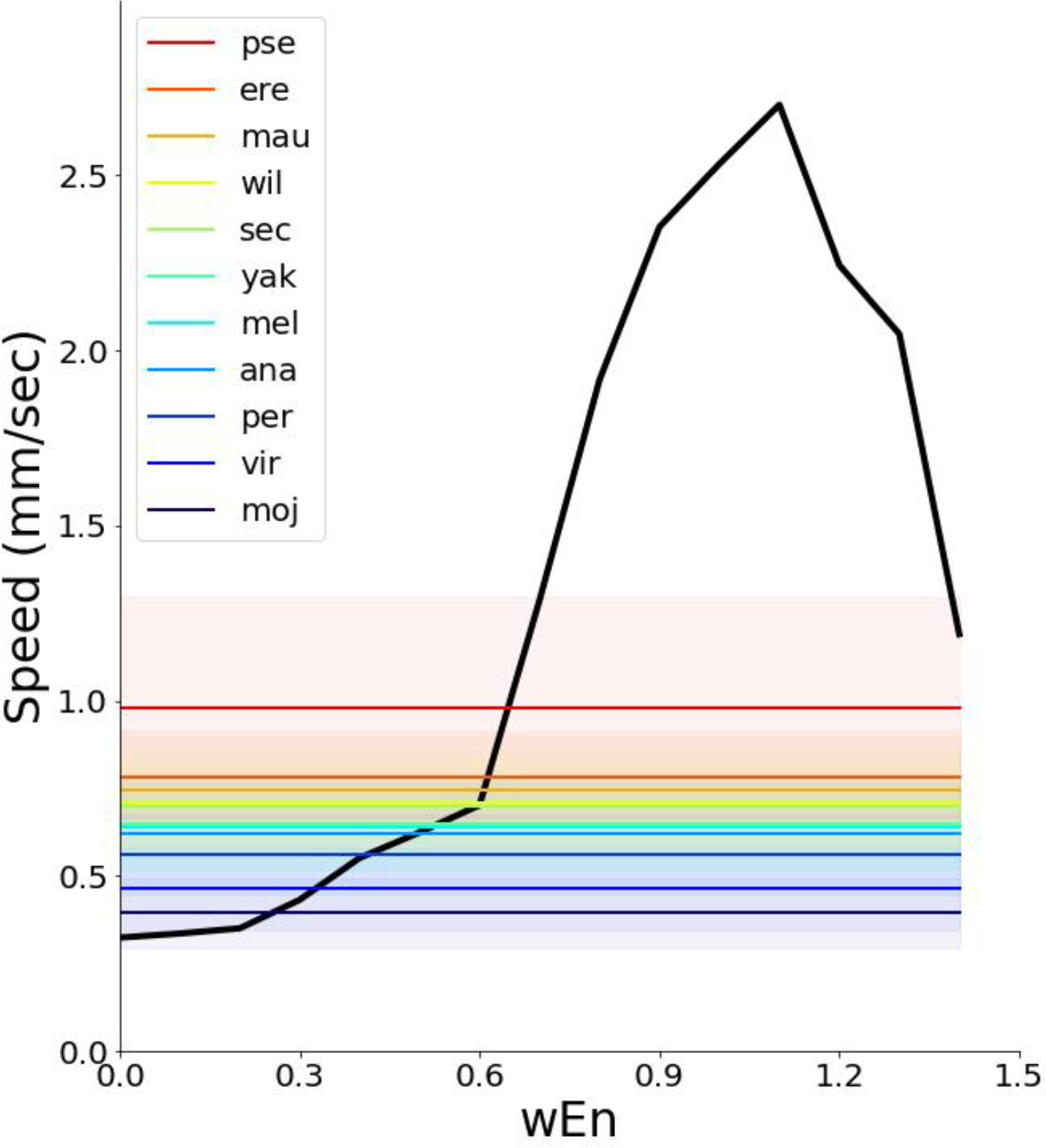
Prediction of the strength of intersegmental connections in sister species in genus *Drosophila.* The black curve shows the relation between the intersegmental coupling *W*_*En*_ and crawling speed obtained by the neuromechanical model based on the measurement of *Drosophila melanogaster* larvae. This plot was obtained by averaging out the weight of proprioceptive sensory feedback *W*_*ES*_ in Figure 9B. The average and standard error mean of speed in species in the genus *Drosophila* are shown by solid lines and shaded area, respectively. The speed data was calculated from data in (57).

## Discussion

In this work, we built a neuromechanical model based on the physical properties of fly larvae. The model successfully described crawling behaviour quantitatively. First, we quantified segmental dynamics in larval crawling and obtained seven parameters that characterize the kinematics of peristalsis (Figures 1 and 2). Then, using a tensile tester, viscoelasticity and tension force of larvae were measured (Figures 3 and 4). These results suggest that the larval body could be modelled as a chain of mass and SLS modules (Figure 4), distinct from the previous modelling for larvae (10,39,41). By incorporating the material properties and optimizing parameters in the neural circuit, our model succeeded in reproducing larval crawling quantitatively (Figure 5). Furthermore, this model could describe the observation in optogenetic studies (Figure 6) and crawling in a low friction condition tested in this study (Figure 7). Perturbation analyses indicated the importance of intersegmental connections in the CNS, contrasting to previous studies (Figures 8 and 9). Based on this observation, we predicted the intersegmental connections in the CNS in sister species in the genus *Drosophila* (Figure 12).

One striking finding from our model contrasts with previous studies: the significant contribution of intersegmental connections in the CNS to crawling speed. Although several interneurons involved in speed control have been reported (32,54–56), previous simulation studies were not consistent with these observations. Our model had larger viscosity and smaller contraction forces (Supplementary Figure 3) than previous parameter regimes, implying that each segment is hard to move. The large weight of neuromuscular connection compensated for this difficulty (*W*_*ME*_) and that of sensory feedback (*W*_*ES*_) (Supplementary Figure 3). With the strong interaction between the CNS and peripheral organs (muscles and proprioceptors), the CNS could play a significant role in controlling crawling speed (Figure 8I). This finding demonstrates the significance of intersegmental connection in the CNS. It also highlights the importance of mechanical properties in the larval body and the dynamics of segments.

We measured the viscoelastic properties of larvae by the stress-relaxation test. It should be noted that there are some limitations in this measurement. First, we obtained these passive properties of the body by measuring unanaesthetised larvae. We analyzed the data without any spontaneous contraction in the stress-relaxation test. However, there remains a possibility that infinitesimal muscular tension is induced by proprioceptive feedback upon stretching. By blocking action potentials in motor neurons while keeping spontaneous vesicle release, it would be able to suppress feedback from proprioception to measure purer viscoelasticity. Second, we approximated the viscoelasticity for the contraction in the range of 0.2 mm by that for the extension in the range of 0.04 mm. In real larval crawling, asymmetricity between extension and contraction and nonlinearity in the viscosity would affect the kinematics. By incorporating these factors, it would be able to reproduce the kinematics more quantitatively.

The model in this study could describe forward crawling, the most frequent behaviour in larvae. Recent neuroscience studies have revealed numerous neural circuit modules involved in distinct aspects of behaviour, including speed control (32), bilateral coordination (35), intrasegmental coordination (36), intersegmental coordination (31), backward crawling (58, 59), turning (60), escaping (61), and sensory input guided navigation (34, 62). By integrating these circuit modules, we would be able to establish a neuromechanical model that reproduces multiple and natural larval behaviour. Furthermore, our model can potentially serve biomimetics. In recent decades, more and more soft robots, endowed with new capabilities relative to the traditional hard ones, have been designed to exhibit complex movements (63–66). Taking *Drosophila* larvae as a prototype, bionic structures can be established with high dexterity to explore the unstructured environments. By harnessing perturbation analyses on the neuromechanical model, the locomotion ability of the soft larval robots could be tuned and optimized for required applications.

## Materials and Methods

### Fly strains

We used the third instar larvae in all the experiments. To quantify segment dynamics, we used *tubP-Gal4*, *UAS-fondue::GFP* (Bloomington #5138, #43646), which labels the segment boundaries (45). In the viscoelasticity measurements, we used a wild type strain *Canton S*. For optogenetic experiments, *OK6-Gal4* (67) and *UAS-ChR2[T159C]::YFP* (31) were used.

### Image acquisition and segment boundary annotation

Third instar larvae of *tubP-Gal4, UAS-fondue::GFP* were gently washed in deionised water to remove residual food from the body surface. Then, individual larvae were placed on a flat agarose stage (1.5% agarose). Behavioural videos were captured by a CCD camera (XCD60, Sony, Japan) mounted on an Olympus stereomicroscope (SZX16, Olympus, Japan). Images were acquired at 30 Hz. We used Fiji software (68) to manually annotate the right and left ends of every segment boundary (from the anterior boundary of the T2 segment to the posterior boundary of the A8 segment. See Figure 1A.) The midpoint of the right and left ends of a boundary was used as the position of the boundary.

### Viscoelasticity measurement

Third instar larvae in the feeding stage were washed with distilled water and dried with paper. Insect pins were inserted into the head and tail. The pin in the head was bent to form a loop to hook to a paper clip that was hung on the hook of the tensile sensing machine, while the pin in the tail was used to fix the body on the PDMS (Polydimethylsiloxane, Sylgard 184, Toray, Japan) silicone block, held by the tong on the SHIMADZU EZ-S platform with a 5 N load cell. The experimental equipment is shown in Figure 4A. The larval body was kept on the vertical axis for measurement. The baseline of force was calibrated by values measured before applying external elongational force. All the experimental procedures were performed at room temperature.

During the stress relaxation tests, we applied a constant strain of 0.4 mm, about 10% of the body length (3.53 ± 0.12 mm, n = 9 larvae), to the larvae. Then, the stress decreased until the plateau was reached after a while, as shown in Figure 4G-4H. To fit the stress relaxation curve to mechanical analogues, we adapted the Maxwell, Kelvin-Voigt, and standard linear solid (SLS) models. These models give the relationship between joint force *F* and displacement Δ*L*. Detailed functions are described as follows:

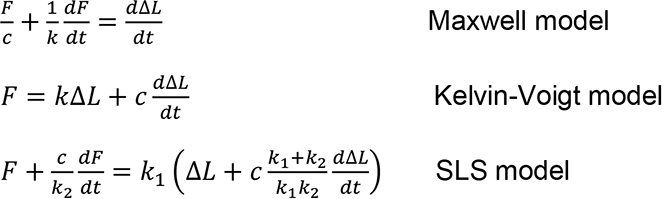

where *k* and *c* are the elastic and damping constants, respectively. The total displacement of 0.4 mm was realised by pulling the larval body with a rate of 1 mm/min. During the stress relaxation experiments, this elongation time (24 sec) is much shorter than that for relaxation (576 sec), and thus the displacement is regarded as the step function *L*(*t*) = *L*_0_*H*(*t* − *t*_0_) and initial force is *F*(0) = 0*N*, where *L*_0_ = 0.4 *mm* and *H*(*t*) is the Heaviside step function. In this case, the corresponding stress relaxation functions in these models are described as follows:

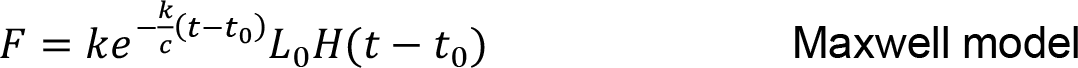

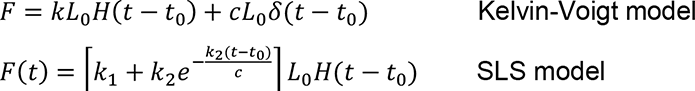

where *t*_0_ ≥ 0. By fitting these curves to the stress-relaxation measurement data, we obtained the spring constants and damping coefficients. We used Python 3.7 for the curve fitting.

### Contraction force measurement with optogenetics

For optogenetic activation of motor neurons, we used the *OK6-GAL4, UAS-ChR2* line (32). The early third-instar larvae were selected and put into ATR (all-trans-retinal) containing yeast paste, the concentration of which was 1 mM. These larvae were reared at 25 °C in the dark for one day (69). Afterwards, the third-instar larvae were prepared to measure tension force as described above (Viscoelasticity measurement section). Blue LED light (455 nm, 5.7 nW/mm^2^, M455L3, ThorLab) was used to stimulate ChR2 expressed in motor neurons, which leads to the contraction of the larval body. In each stimulation, the blue light was applied for two seconds followed by a no-illumination interval of two seconds, and the force induced was monitored. Eighteen larvae were used in the measurement, and each measurement took two to five minutes. The optogenetically induced forces were measured by the differences between forces during illumination and no illumination (Figure 3).

### Larval crawling in a low-friction environment

We used a high concentration sugar solution (66% w/w sucrose) and floated third- instar larvae in it. The genotype we used was *R70C01-Gal4, UAS-CD4::GFP*, which allowed us to mark the segmental boundary from T2 to A8 (Figure 1A and 1B). The locomotion was recorded via SZX16 fluorescent microscope (Olympus, Japan) with a 1.25x object lens at 30 frames/sec, and its trajectory was measured by Fiji (68).

### Modelling

#### Body-substrate mechanics and modelling

As we mentioned before, the larval crawling stride consists of a piston phase and a wave phase (26). The piston phase constrains the larval body length to be almost constant, while the wave phase generates the propagation wave repetitively. To make it simple, we modelled the whole body of larvae as a chain of eleven segments.

We assumed that the model framework possesses a constant length and its segments are entirely repetitive based on the observation in Figures 1 and 2. Then the body is modelled as a chain of the SLS units in series (Supplementary Figure 1). Muscle groups are modelled as the tension actuator to accept efferent control from the neural circuit (Figure 5B). The tension *F*_*m*_ is the contraction force to counteract the effect of body viscoelasticity and friction. The mechanics are described based on Newton’s second law as follows.

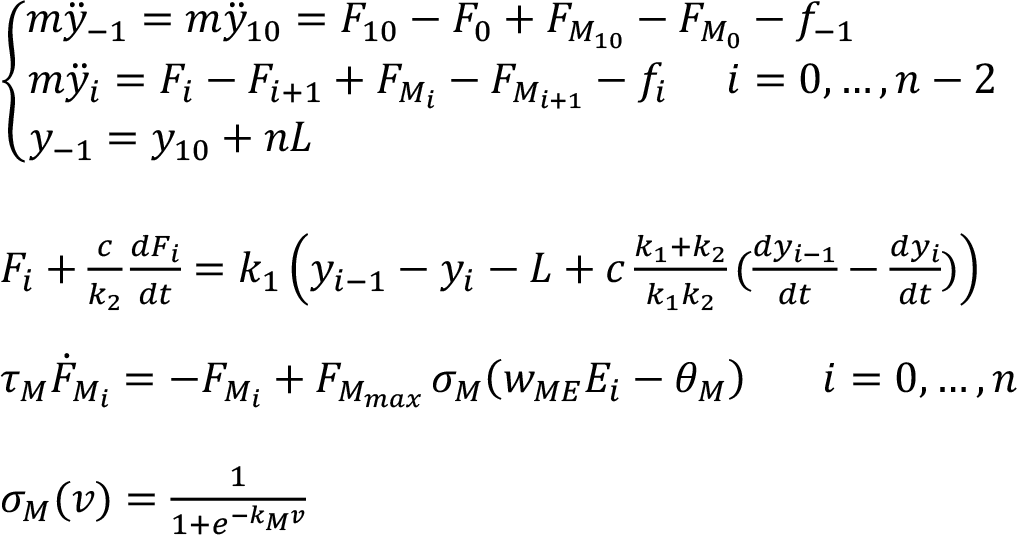

where *y*_*i*_, is the segmental boundary position (Figure 1B and Supplementary Figure 1), *m* is mass, *F* is viscoelastic force, *F*_*M*_ is tension force, τ_*M*_ is time relaxation constant for tension, *W*_*ME*_ is connection weight from excitatory unit to tension actuator, θ_*M*_ is the threshold for tension, and *k*_*m*_ is the gain in function σ_*M*_(*v*), and *n* is the total number of segments, equal to 11. *y*_−1_ is the position of the head and *y*_10_ is the position of the tail. To model the piston phase, where the head and tail move concurrently, the posterior and anterior ends share the same velocity during crawling, as *y*_−1_ = *y*_10_ which is inferred from the third equation above.

Each mass block is dependent on joint force from SLS modules, tensions, and friction. The joint force from the SLS unit is affected by the segmental displacement, and the tension force is regulated by the sigmoid function with gain *k*_*t*_. The periodic tension force, modelled by the sigmoid function of excitatory inputs with gain *k*_M_, is indispensable to sustain the propagation of contraction wave, verified by the experimental results (53).

As for the friction force, it exists during the interaction of the mechanical body with the substrate. The transition process from static to dynamic friction is not modelled in this work. Instead, the ratio of forward to backward friction is introduced to the model as directional asymmetric, considering the anterior-posterior polarity in denticle bands. Since the backward friction is larger than forward friction, the ratio is more than one. Friction on the segmental boundary is represented as:

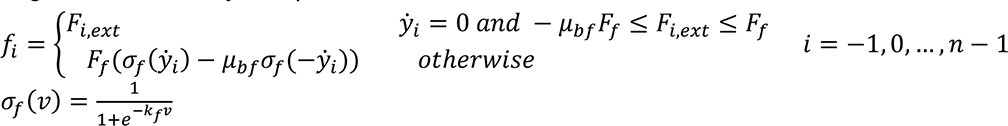

where *f*, *F*_*f*_, μ_*bf*_(_*bf*_ > 1) individually represent the friction, forward friction and ratio for forward-backward friction, and *F*_*i,ext*_ refers to the joint force of all the other forces. When the mass block is still and *F*_*i,ext*_ does not exceed the range of forward-backward friction, the friction force *f*_*i*_ should be equal to *F*_*i,ext*_. Otherwise, the friction is either the forward friction *F*_*f*_ or the backward one μ_*bf*_*F*_*f*_.

### Neuromuscular dynamics and modelling

The framework for the neural circuit is depicted in Figure 5B. The neural circuit model is based on the model in Phelevan *et al*. (2016). Neural dynamics is realized by the activities of excitatory and inhibitory populations of neurons in each segment under the Wilson-Cowan model (70, 71). In *Drosophila* larvae, most proprioceptive neurons were active when the segment was contracted (43). The feedback from the sensory receptor provides signals for the inhibitory neuron within the same segment and excitatory neuron in the anterior segment. In this case, it can promote both local relaxation and forward propagation waves. This connectivity is consistent with the “mission-accomplished” model (44).

Under these assumptions, neural dynamics of neural circuits are depicted as follows:

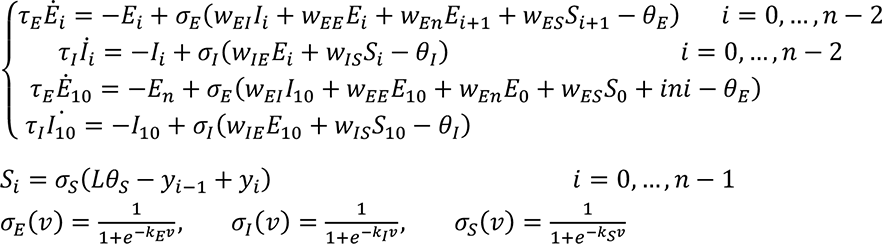

where *E* and *I* are the mean firing rates of the excitatory and inhibitory population, τ_*E*_ and τ_*1*_ are the relaxation time constant of the excitatory and inhibitory units, respectively, *W*_*a,b*_ is their synaptic connection weight from population *b* to population *a*, *θ*_*E*_ and *θ*_*I*_ are the activation threshold of the excitatory and inhibitory units, *k*_*E*_, *k*_*I*_, and *k*_*S*_ are the sigmoid gain of excitatory, inhibitory, and sensory units, respectively, and *S* is the feedback strength from the sensory receptor. The circuit includes excitatory connection via *W*_*IE*_, *W*_*ME*_ and inhibitory connection via *W*_*ES*_, *W*_*EI*_ . When the segmental length becomes smaller than the threshold *Lθ*_*s*_, the sensory unit starts to be activated. To trigger the initial crawling, we introduced an external stimulus *ini*, a rectangular pulse (for 5 ms), on the excitatory population in the posterior terminal.

The values of the parameters are listed in Supplementary Figure 3. All the simulation work is performed using stiff solver ode15s in MATLAB R2020a.

## Acknowledgement

We are grateful to L.C. Griffith, H. Aberle, Bloomington Drosophila Stock Center, and the Kyoto Stock Center for fly stocks. We thank C. Pehlevan for his critical reading of the paper.

## Supporting Information

**Supplementary Figure 1.**
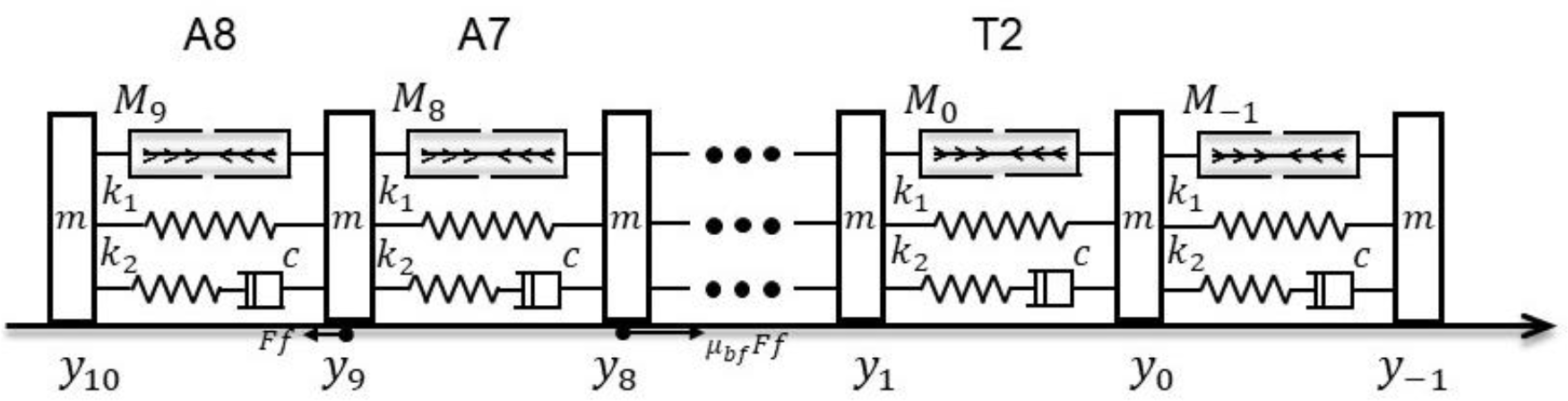
A physical model with eleven segments. Based on the estimation of the number of segments, the fly larva was modelled by eleven segments. We assumed that *y*_10_ and *y*_−1_ were physically coupled. In the simulation results in this paper, segmental boundaries from *y*_10_ to *y*_0_ or segments from A8 to T2 were shown.

**Supplementary Figure 2.**
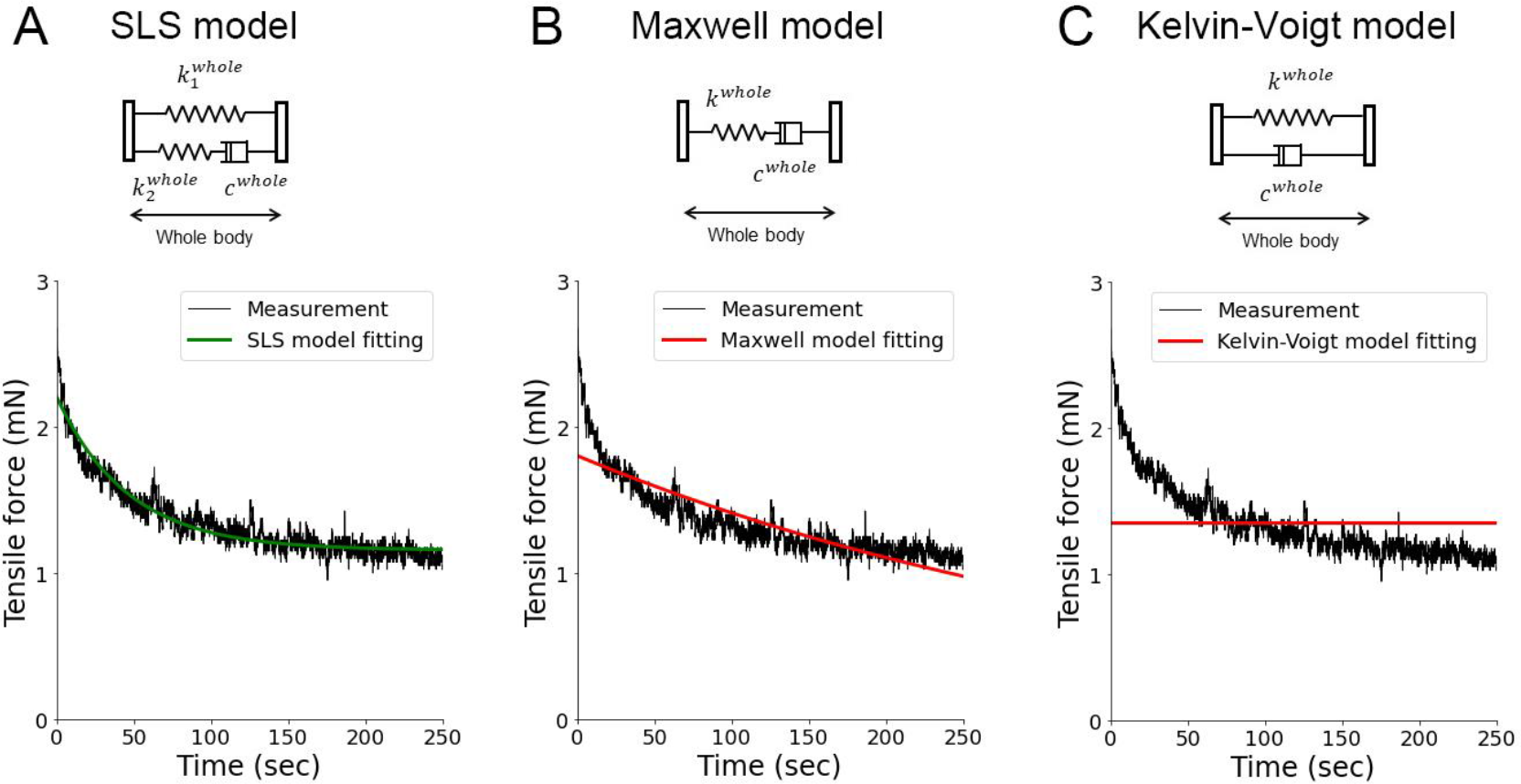
Fitting stress-relaxation test data by three viscoelastic models. The same stress-relaxation test data was fitted with the SLS model (A), Maxwell model (B), and Kelvin-Voigt model (C).

**Supplementary Figure 3.**
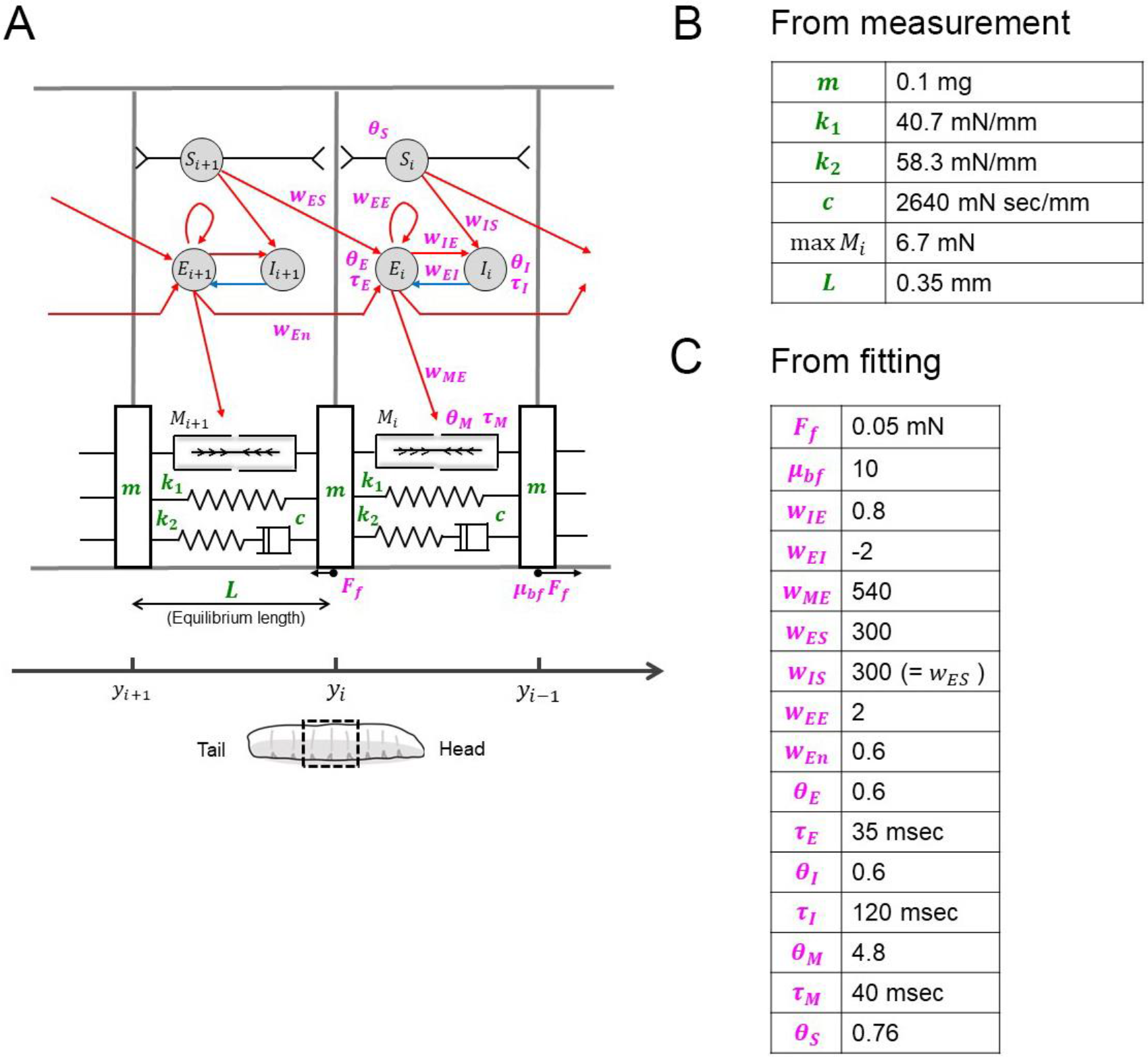
Our physical model for larval crawling and its parameters. Schematic of the physical model for larval crawling. (B) List of parameters whose values were obtained by measurement using fly larvae. (C) List of parameters whose values were obtained by fitting to reproduce larval crawling by the simulation.

**Supplementary Figure 4.**
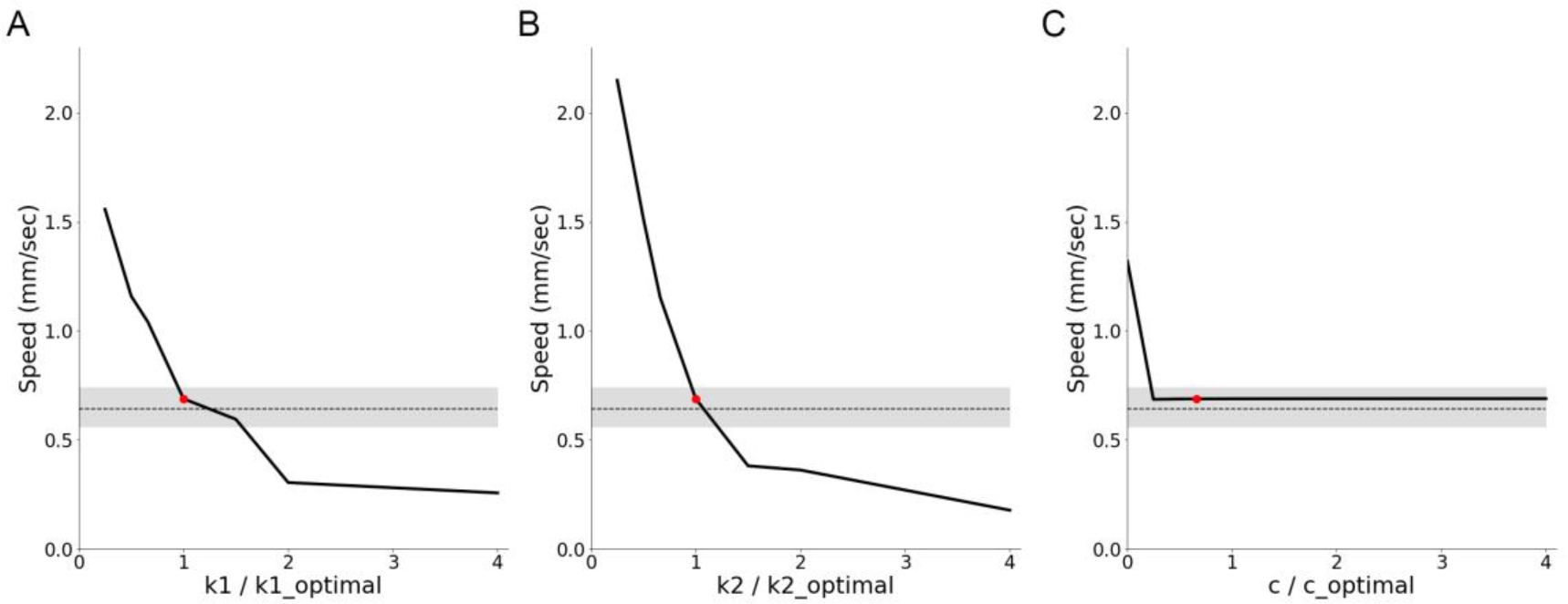
Perturbation analysis on viscoelasticity Plots of speed as viscoelasticity was perturbed. (A) Perturbation of the spring constant *k*_1_ Perturbation of the spring constant *k*_2_. (C) Perturbation of the dumping coefficient *c*. The horizontal axes were normalized by the optimized values (*k*_1__*optimal*, *k*_2__*optimal*), respectively. Grey shaded regions show the range of speed observed in the experiment with third-instar larvae. Red dots indicate the optimized simulation condition. The minimum value in the horizontal axis in the plot of (C) equals 1/400, which corresponds to the value in Pehlevan *et al.* (2016).

**Supplementary Figure 5.**
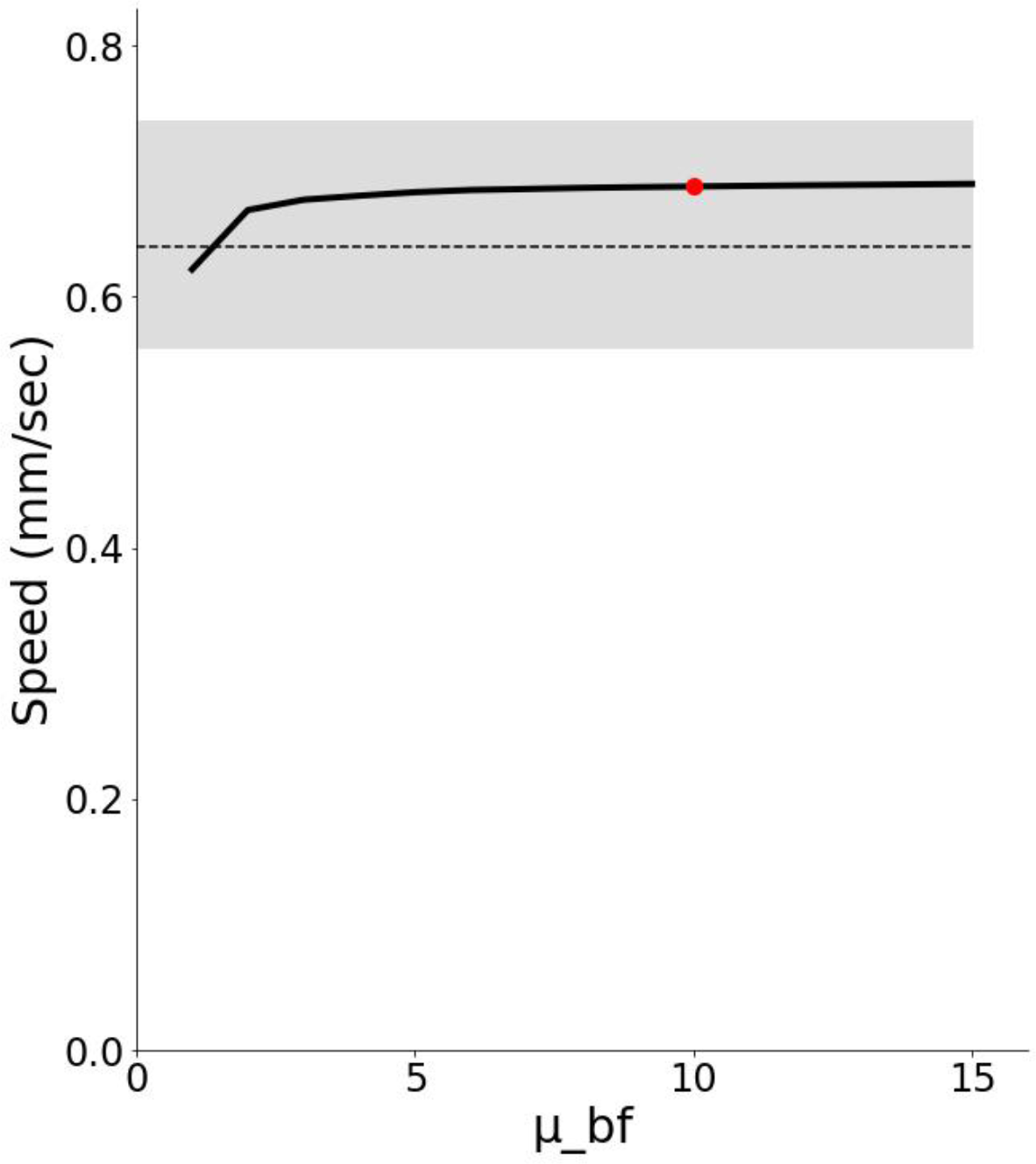
Perturbation analysis on the asymmetricity in friction Plots of speed as the friction asymmetricity was perturbed. The grey shaded region shows the range of speed observed in the experiment with third-instar larvae. The red dot indicates the optimized simulation condition.

**Supplementary Figure 6.**
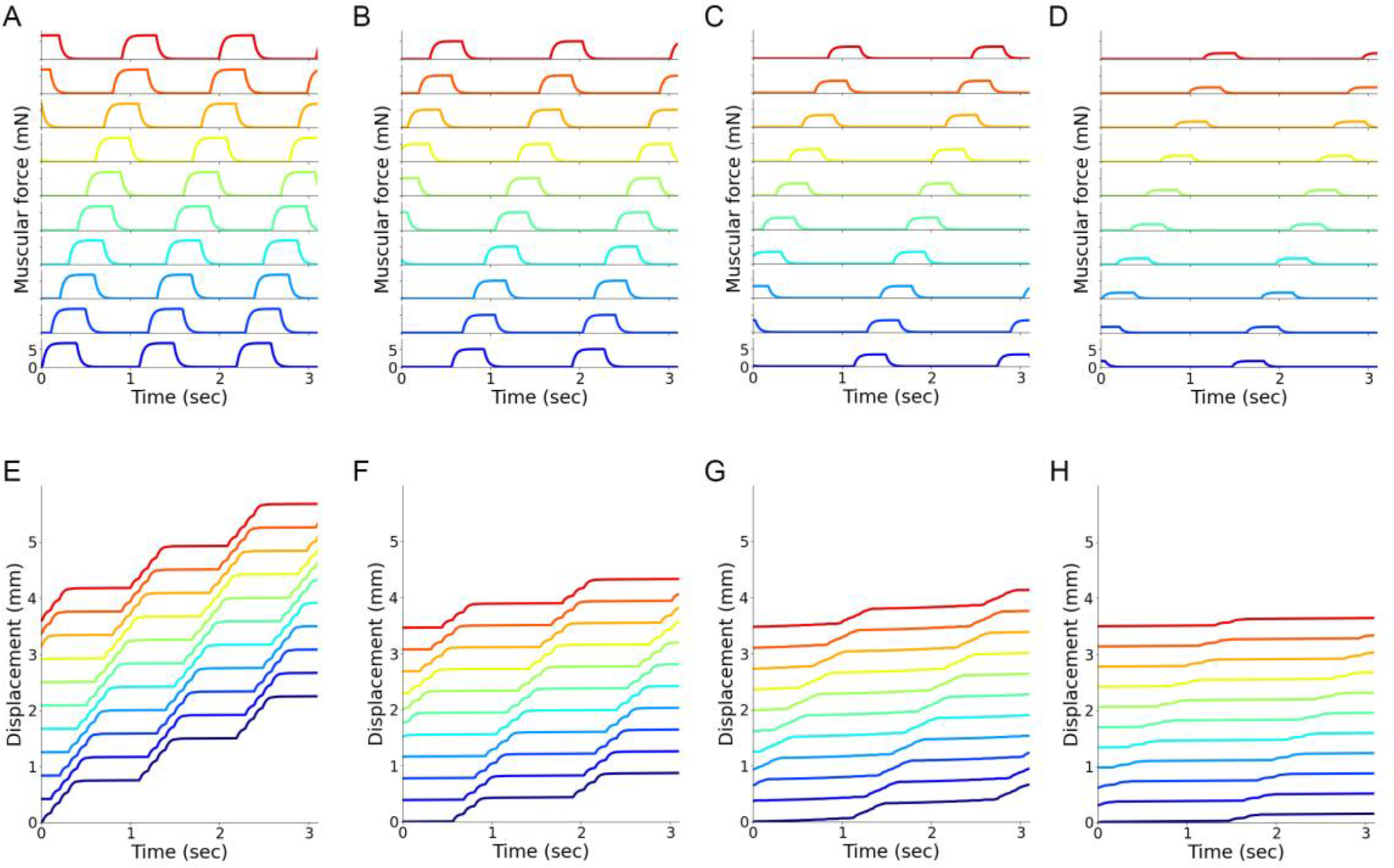
Perturbation analysis on the maximum muscular force. Muscular force and segment displacement when the maximum muscular contraction force was perturbed. (A-D) Muscular forces in all the segments. Line colours correspond to those in Figure 1B and 1C. (E-H) Displacement of the segmental boundaries. Line colours correspond to those in Figure 1E and 1F. The ratio of the maximum contraction force to the optimized force: A and E, 100%; B and F, 75%; C and G, 50%; D and H, 25%.

**Supplementary Figure 7.**
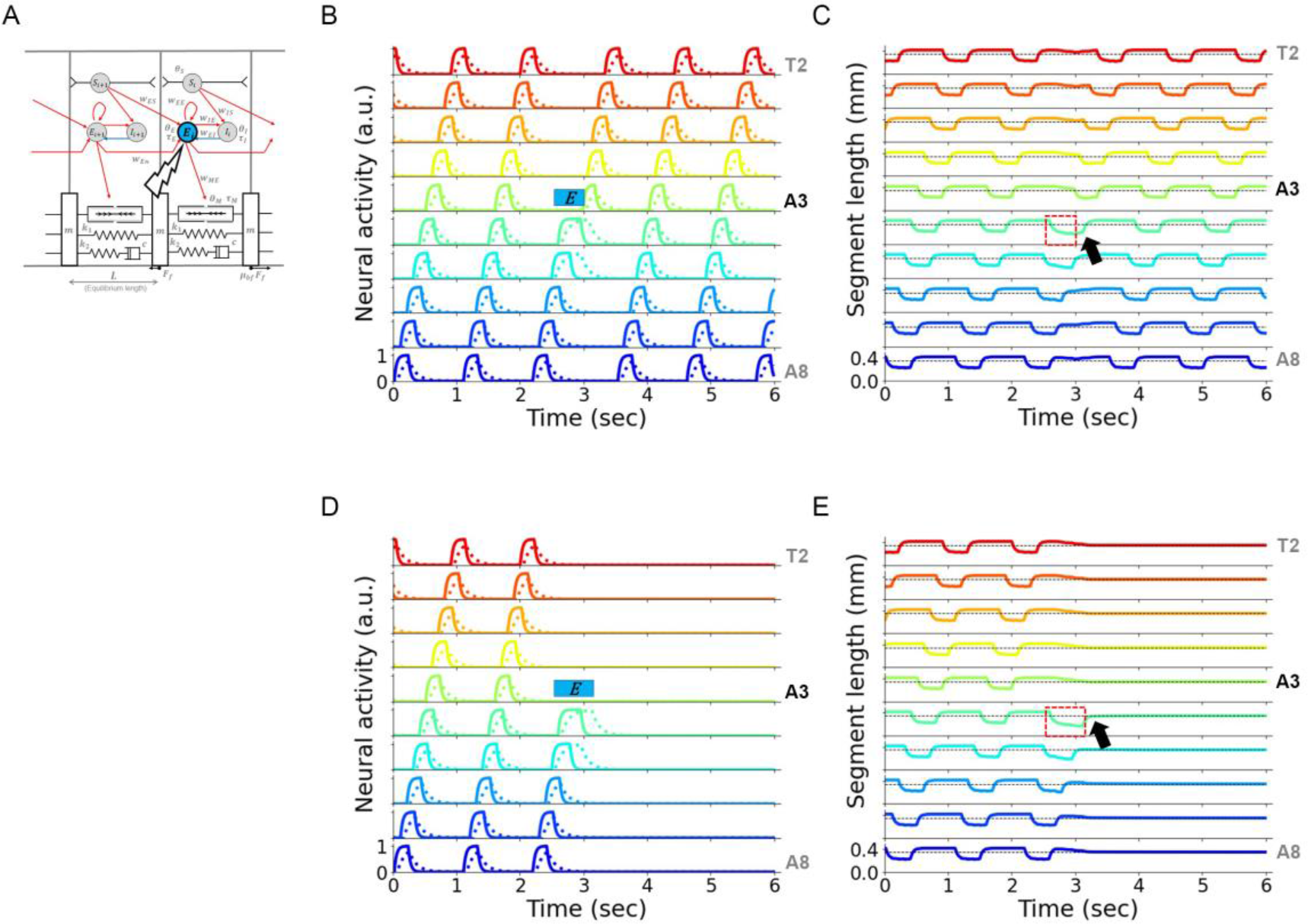
Different durations of optogenetic stimulation in the simulation exhibit distinct phenotypes in the propagation of waves. (A) Schematic of optogenetic silencing of excitatory neurons in a single segment shown by a blue disk. (B) Neural activity with the optogenetic silencing of excitatory neurons. Excitatory neurons in the A3 segment were silenced for 0.5 seconds, marked by a blue bar. Waves were resumed just after the optogenetic silencing was removed. (C) Traces of segment length in (B). The dotted box shows the segment length of the posterior neighbouring segment during optogenetic stimulation. The arrow indicates that the neighbouring segment was contracted just after the optogenetic stimulus was removed. (D) Neural activity with the optogenetic silencing of excitatory neurons for 0.7 seconds. Excitatory neurons in the A3 segment were silenced as marked by a blue bar. Waves were arrested even after the optogenetic silencing was removed. (E) Traces of segment length in (D). The dotted box indicates the segment length of the posterior neighbouring segment during optogenetic stimulation. The arrow indicates that the neighbouring segment was returned to the equilibrium length just after the optogenetic stimulus was removed.

**Supplementary Figure 8.**
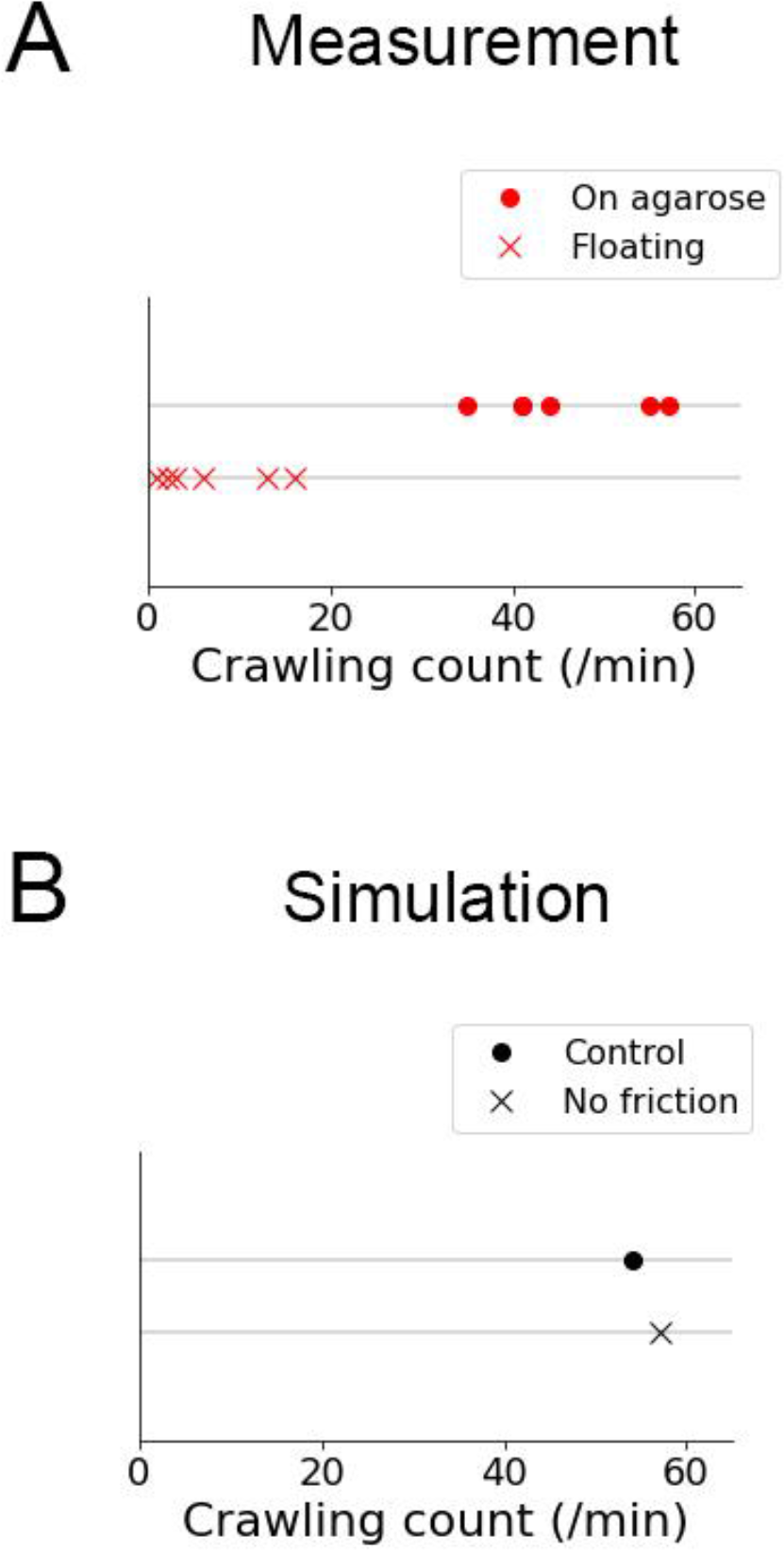
Frequency of crawling in a low friction condition. (A) Comparison of crawling frequency between larvae on agarose and floating. (B) Comparison of crawling frequency in simulation between in the optimized condition (Control) and the absence of friction (No friction).

## References

1. Grillner S, El Manira A. Current principles of motor control, with special reference to vertebrate locomotion. Physiol Rev. 2020;100(1):271–320.

2. Ijspeert AJ. Central pattern generators for locomotion control in animals and robots: A review. Neural Networks. 2008;21(4):642–53.

3. Marder E, Calabrese RL. Principles of rhythmic motor pattern generation. Physiol Rev. 1996;76(3):687–717.

4. Marder E, Bucher D. Central pattern generators and the control of rhythmic movements. Curr Biol. 2001;11(23):R986–96.

5. Minassian K, Hofstoetter US, Dzeladini F, Guertin PA, Ijspeert A. The Human Central Pattern Generator for Locomotion: Does It Exist and Contribute to Walking? Neuroscientist. 2017;23(6):649–63.

6. Gomez-Marin A, Ghazanfar AA. The Life of Behavior. Neuron. 2019;104(1):25–36.

7. Scheffer LK, Meinertzhagen IA. A connectome is not enough – What is still needed to understand the brain of Drosophila? J Exp Biol. 2021;224(21).

8. Miller LA, Goldman DI, Hedrick TL, Tytell ED, Wang ZJ, Yen J, et al. Using computational and mechanical models to study animal locomotion. Integr Comp Biol. 2012;52(5):553–75.

9. Tytell ED, Holmes P, Cohen AH. Spikes alone do not behavior make: Why neuroscience needs biomechanics. Curr Opin Neurobiol. 2011;21(5):816–22.

10. Pehlevan C, Paoletti P, Mahadevan L. Integrative neuromechanics of crawling in D. Melanogaster Larvae. Elife. 2016;5(2016JULY):1–23.

11. Boyle JH, Berri S, Cohen N. Gait modulation in C. elegans: An integrated neuromechanical model. Front Comput Neurosci. 2012;6(FEBRUARY 2012):1–15.

12. Ekeberg Ö, Grillner S. Simulations of neuromuscular control in lamprey swimming. Philos Trans R Soc B Biol Sci. 1999;354(1385):895–902.

13. Russo A Di, Stanev D, Armand S, Ijspeert A. Sensory modulation of gait characteristics in human locomotion: A neuromusculoskeletal modeling study. Vol. 17, PLoS Computational Biology. 2021. 1–33 p.

14. Ijspeert AJ, Crespi A, Cabelguen JM. Simulation and robotics studies of salamander locomotion. Neuroinformatics. 2005;3(3):171–95.

15. Nishikawa K, Biewener AA, Aerts P, Ahn AN, Chiel HJ, Daley MA, et al. Neuromechanics: An integrative approach for understanding motor control. Integr Comp Biol. 2007;47(1):16–54.

16. Elder HY. Peristaltic mechanisms. In: Aspects of Animal Movement [Society for Experimental Biology Seminar Series, vol5]. Cambridge UP; 1980. p. 71–92.

17. Alexander RM. Principles of Animal Locomotion. Princeton University Press; 2006. 384 p.

18. Lin HT, Dorfmann AL, Trimmer BA. Soft-cuticle biomechanics: A constitutive model of anisotropy for caterpillar integument. J Theor Biol. 2009;256(3):447–57.

19. Dorfmann AL, Woods WA, Trimmer BA. Muscle performance in a soft-bodied terrestrial crawler: Constitutive modelling of strain-rate dependency. J R Soc Interface. 2008;5(20):349–62.

20. Paterson BA, Marko Anikin I, Krans JL. Hysteresis in the production of force by larval Dipteran muscle. J Exp Biol. 2010;213(14):2483–93.

21. Backholm M, Ryu WS, Dalnoki-Veress K. Viscoelastic properties of the nematode Caenorhabditis elegans, a self-similar, shear-thinning worm. Proc Natl Acad Sci U S A. 2013;110(12):4528–33.

22. Gilpin W, Uppaluri S, Brangwynne CP. Worms under pressure: Bulk mechanical properties of C. elegans are independent of the cuticle. Biophys J. 2015;108(8):1887–98.

23. Rabets Y, Backholm M, Dalnoki-Veress K, Ryu WS. Direct measurements of drag forces in c. elegans crawling locomotion. Biophys J. 2014;107(8):1980–7.

24. Berrigan D, Pepin DJ. How maggots move: Allometry and kinematics of crawling in larval Diptera. J Insect Physiol. 1995;41(4):329–37.

25. Green CH, Burnet B, Connolly KJ. Organization and patterns of inter- and intraspecific variation in the behaviour of Drosophila larvae. Anim Behav. 1983;31(1):282–91.

26. Heckscher ES, Lockery SR, Doe CQ. Characterization of Drosophila larval crawling at the level of organism, segment, and somatic body wall musculature. J Neurosci. 2012;32(36):12460–71.

27. Lahiri S, Shen K, Klein M, Tang A, Kane E, Gershow M, et al. Two alternating motor programs drive navigation in Drosophila larva. PLoS One. 2011;6(8).

28. Fox LE, Soll DR, Wu CF. Coordination and modulation of locomotion pattern generators in Drosophila larvae: Effects of altered biogenic amine levels by the tyramine β hydroxlyase mutation. J Neurosci. 2006;26(5):1486–98.

29. Hunter I, Coulson B, Zarin AA, Baines RA. The Drosophila Larval Locomotor Circuit Provides a Model to Understand Neural Circuit Development and Function. Front Neural Circuits. 2021;15(July):1–12.

30. Gowda SBM, Salim S, Mohammad F. Anatomy and neural pathways modulating distinct locomotor behaviors in Drosophila larva. Biology (Basel). 2021;10(2):1–30.

31. Kohsaka H, Zwart MF, Fushiki A, Fetter RD, Truman JW, Cardona A, et al. Regulation of forward and backward locomotion through intersegmental feedback circuits in Drosophila larvae. Nat Commun. 2019;10(1):2654.

32. Kohsaka H, Takasu E, Morimoto T, Nose A. A group of segmental premotor interneurons regulates the speed of axial locomotion in Drosophila larvae. Curr Biol. 2014;24(22):2632–42.

33. Clark MQ, Zarin AA, Carreira-Rosario A, Doe CQ. Neural circuits driving larval locomotion in Drosophila. Neural Dev. 2018;13(1):1–10.

34. Kohsaka H, Guertin PA, Nose A. Neural Circuits Underlying Fly Larval Locomotion. Curr Pharm Des. 2017;23(12):1722–33.

35. Heckscher ES, Zarin AA, Faumont S, Clark MQ, Manning L, Fushiki A, et al. Even- Skipped+ Interneurons Are Core Components of a Sensorimotor Circuit that Maintains Left-Right Symmetric Muscle Contraction Amplitude. Neuron. 2015;88(2):314–29.

36. Zwart MF, Pulver SR, Truman JW, Fushiki A, Fetter RD, Cardona A, et al. Selective Inhibition Mediates the Sequential Recruitment of Motor Pools. Neuron. 2016;91(3):615– 28.

37. Zarin AA, Mark B, Cardona A, Litwin-Kumar A, Doe CQ. A multilayer circuit architecture for the generation of distinct locomotor behaviors in Drosophila. Elife. 2019;8:1–34.

38. Gjorgjieva J, Berni J, Evers JF, Eglen SJ. Neural circuits for peristaltic wave propagation in crawling Drosophila larvae: analysis and modeling. Front Comput Neurosci. 2013;7:1– 19.

39. Ross D, Lagogiannis K, Webb B. A model of larval biomechanics reveals exploitable passive properties for efficient locomotion. Lect Notes Comput Sci (including Subser Lect Notes Artif Intell Lect Notes Bioinformatics). 2015;9222:1–12.

40. Loveless J, Lagogiannis K, Webb B. Modelling the neuromechanics of exploration and taxis in larval Drosophila. PLoS Comput Biol. 2019;1–33.

41. Paoletti P, Mahadevan L. A proprioceptive neuromechanical theory of crawling. Proc R Soc B Biol Sci. 2014;281(1790).

42. Berni J. Genetic dissection of a regionally differentiated network for exploratory behavior in drosophila larvae. Curr Biol. 2015;25(10):1319–26.

43. Vaadia RD, Li W, Voleti V, Singhania A, Hillman EMC, Grueber WB. Characterization of Proprioceptive System Dynamics in Behaving Drosophila Larvae Using High-Speed Volumetric Microscopy. Curr Biol. 2019;0(0):1–10.

44. Hughes CL, Thomas JB. A sensory feedback circuit coordinates muscle activity in Drosophila. Mol Cell Neurosci. 2007;35(2):383–96.

45. Green N, Odell N, Zych M, Clark C, Wang ZH, Biersmith B, et al. A common suite of coagulation proteins function in drosophila muscle attachment. Genetics. 2016;204(3):1075–87.

46. Bate M. The Development of Drosophila melanogaster. Cold Spring Harbor Laboratory Press; 1993.

47. Pulver SR, Pashkovski SL, Hornstein NJ, Garrity PA, Griffith LC. Temporal dynamics of neuronal activation by channelrhodopsin-2 and TRPA1 determine behavioral output in Drosophila larvae. J Neurophysiol. 2009;101(6):3075–88.

48. Sanyal S. Genomic mapping and expression patterns of C380, OK6 and D42 enhancer trap lines in the larval nervous system of Drosophila. Gene Expr Patterns. 2009;9(5):371–80.

49. Trueman ER (Edwin R. The locomotion of soft-bodied animals. Edward Arnold; 1975. 200 p.

50. Quillin K. Ontogenetic scaling of hydrostatic skeletons: geometric, static stress and dynamic stress scaling of the earthworm lumbricus terrestris. J Exp Biol. 1998;201(12).

51. Quillin KJ. Kinematic scaling of locomotion by hydrostatic animals: ontogeny of peristaltic crawling by the earthworm lumbricus terrestris. J Exp Biol. 1999;202(6).

52. Banks HT, Hu S, Kenz ZR. A brief review of elasticity and viscoelasticity for solids. Adv Appl Math Mech. 2011;3(1):1–51.

53. Inada K, Kohsaka H, Takasu E, Matsunaga T, Nose A. Optical dissection of neural circuits responsible for drosophila larval locomotion with Halorhodopsin. PLoS One. 2011 Dec;6(12):e29019.

54. Fushiki A, Zwart MF, Kohsaka H, Fetter RD, Cardona A, Nose A. A circuit mechanism for the propagation of waves of muscle contraction in Drosophila. Elife. 2016;5:1–23.

55. Yoshikawa S, Long H, Thomas JB. A subset of interneurons required for Drosophila larval locomotion. Mol Cell Neurosci. 2016;70:22–9.

56. Babski H, Surel C, Yoshikawa S, Valmier J, Thomas JB, Carroll P, et al. A GABAergic Maf-expressing interneuron subset regulates the speed of locomotion in Drosophila. Nat Commun. 2019;10(4796):1–17.

57. Matsuo Y, Nose A, Kohsaka H. Interspecies variation of larval locomotion kinematics in the genus Drosophila and its relation to habitat temperature. BMC Biol. 2021;19(1):1–21.

58. Hiramoto A, Jonaitis J, Niki S, Kohsaka H, Fetter RD, Cardona A, et al. Regulation of coordinated muscular relaxation in Drosophila larvae by a pattern-regulating intersegmental circuit. Nat Commun. 2021;12(1):1–14.

59. Carreira-Rosario A, Zarin AA, Clark MQ, Manning L, Fetter RD, Cardona A, et al. MDN brain descending neurons coordinately activate backward and inhibit forward locomotion. Elife. 2018;7:1–28.

60. Tastekin I, Riedl J, Schilling-Kurz V, Gomez-Marin A, Truman JW, Louis M. Role of the subesophageal zone in sensorimotor control of orientation in drosophila larva. Curr Biol. 2015;25(11):1448–60.

61. Ohyama T, Schneider-Mizell CM, Fetter RD, Aleman JV, Franconville R, Rivera-Alba M, et al. A multilevel multimodal circuit enhances action selection in Drosophila. Nature. 2015;520(7549):633–9.

62. Louis M. Mini-brain computations converting dynamic olfactory inputs into orientation behavior. Curr Opin Neurobiol. 2020;64:1–9.

63. Kim S, Laschi C, Trimmer B. Soft robotics: A bioinspired evolution in robotics. Trends Biotechnol. 2013;31(5):287–94.

64. Aguilar J, Zhang T, Qian F, Kingsbury M, McInroe B, Mazouchova N, et al. A review on locomotion robophysics: The study of movement at the intersection of robotics, soft matter and dynamical systems. Reports Prog Phys. 2016;79(11).

65. Trivedi D, Rahn CD, Kier WM, Walker ID. Soft robotics: Biological inspiration, state of the art, and future research. Appl Bionics Biomech. 2008;5(3):99–117.

66. Corucci F, Cheney N, Giorgio-Serchi F, Bongard J, Laschi C. Evolving soft locomotion in aquatic and terrestrial environments: Effects of material properties and environmental transitions. Soft Robot. 2018;5(4):475–93.

67. Aberle H, Haghighi AP, Fetter RD, McCabe BD, Magalhães TR, Goodman CS. Wishful thinking encodes a BMP type II receptor that regulates synaptic growth in Drosophila. Neuron. 2002;33(4):545–58.

68. Schindelin J, Arganda-Carreras I, Frise E, Kaynig V, Longair M, Pietzsch T, et al. Fiji: An open-source platform for biological-image analysis. Vol. 9, Nature Methods. Nature Publishing Group; 2012. p. 676–82.

69. Matsunaga T, Fushiki A, Nose A, Kohsaka H. Optogenetic perturbation of neural activity with laser illumination in semi-intact drosophila larvae in motion. J Vis Exp. 2013;(77):1– 5.

70. Wilson HR, Cowan JD. Excitatory and Inhibitory Interactions in Localized Populations of Model Neurons. Vol. 12, Biophysical Journal. 1972.

71. Negahbani E, Steyn-Ross DA, Steyn-Ross ML, Wilson MT, Sleigh JW. Noise-Induced Precursors of State Transitions in the Stochastic Wilson–Cowan Model. J Math Neurosci. 2015 Dec;5(1):1–27.

